# Plant-pollinator specialization: Origin and measurement of curvature

**DOI:** 10.1101/2021.10.05.429567

**Authors:** Mannfred M.A. Boehm, Jill E. Jankowski, Quentin C.B. Cronk

## Abstract

A feature of biodiversity is the abundance of curves displayed by organs and organisms. Curvature is a widespread, convergent trait that has important ecological and evolutionary implications. In pollination ecology, the curvature of flowers and pollinator mouthparts (e.g. hummingbird bills) along the dorsiventral plane has been associated with specialization, competition, and species co-existence. Six methods have historically been used to measure curvature in pollination systems; we provide a solution to this inconsistency by defining curvature using well-established concepts from differential geometry. Intuitively, curvature is the degree to which a line is not straight, but more formally, it is the rate at which the tangent of a curve changes direction with respect to arc length. Here, we establish a protocol wherein a line is fitted against landmarks placed on an image of a curved organ or organism, then curvature is computed at many points along the fitted line and the sum taken. The protocol is demonstrated by studying the development of nectar spur curvature in the flowering plant genus *Epimedium* (Berberidaceae). By clarifying the definition of curvature, our aim is to make the language of comparative morphology more precise and broadly applicable to capture other curved structures in nature.

## The ecology of flower-pollinator curvature

> *“We are beginning to understand why some hummingbird bills are long, whereas others are short, and why some hummingbird flowers are wide, whereas others are narrow. Now, why are bills of some hummingbirds and the tubes of the flowers they visit curved?”*
>
> — – Temeles (1996).

At the center of plant-pollinator diversification is a remarkable variety of floral form. The notion that plant communities experience selection to reduce interspecific mating (“floral isolation,” Grant, 1949) points to the importance of floral diversity in initiating and reinforcing reproductive isolation (Armbruster and Muchhala, 2009). For example, patterns of character displacement in sympatric *Centropogon* C.Presl (Campanulaceae) suggest that competition for pollinators led to the divergence of floral traits associated with bat and hummingbird pollination (Lagomarsino and Muchhala, 2019). In the case of South African *Lapeirousia* Pourr. (Iridaceae), geographic variation in floral tube length has subsequently initiated reproductive isolation between morphs with short and long corolla tubes, despite sharing the same fly pollinator (Minnaar et al., 2019). While patterns of plant-pollinator evolution point to both contemporaneous and asymmetrical co-adaptation (Cardinal and Danforth, 2013; Tripp and McDade, 2013), floral morphology can both be the cause and result of plant-pollinator diversification (Kay and Sargent, 2009; Niet and Johnson, 2012; Ollerton, 2017).

Flower-pollinator curvature as viewed from the side (dorsiventral plane) has been a trait of special interest throughout the post-Darwin era of pollination ecology. However, some floral curvature has origins that may precede any particular ecological function. Instead, curved flower parts (e.g. nectar spurs) might develop within buds when constrained for space. That is, during bud development nectar spurs elongate and curve when met with resistance from the enclosing bud tissue. Following bud opening flower parts may straighten but retain some degree of curvature at maturity. While curvature may thus originate from a developmental constraint or lack of selection for straightness, there can be positive selection for curvature when it increases the complexity of nectar extraction and mechanical interaction between the anthers and the body of the pollinator (Young, 2008; Collins, 2008).

One consequence of floral curvature is pollinator partitioning. In making pollinator observations of the Cape flora, Scott-Elliott (1890) noticed that the flowers of *Leonotis ocymifolia* (Burm.f.) Iwarsson (Lamiaceae) visited by sunbirds (Nectariniidae) were “curved with the same curvature as that of the bird’s beak.” (p. 272). Robertson (1889) insightfully notes that the curved nectar spur of *Viola* L. (Violaceae) “serves to limit the insect visits much more than the mere length of the spur.” (p. 172). Stiles (1975) first posited that Neotropical *Heliconia* L. (Zingerberales) partition hummingbird (Trochilidae) visitation by flower and bill curvature, and that specialization by curve-billed hummingbirds allow co-existence within this species-rich clade. Subsequent research supports this hypothesis (Maglianesi et al., 2014): along the slopes of the Central Cordillera of Costa Rica, the degree of flower and hummingbird bill curvature is proportional to plant-pollinator interaction strength (sensu Dehling et al., 2014) and extent of specialization (*sensu* Blüthgen et al., 2006). In addition to interspecific partitioning of resources, many hummingbirds exhibit sexual dimorphism in bill shape: females have greater bill curvature than males and forage for nectar from curved flowers (Temeles et al., 2005, 2010). Explanations for this pattern range from reduced competition for nectar resources between sexes (Paton and Collins, 1989; Temeles et al., 2010) to selection for mechanically superior bill shapes during male-male competition for territory (Rico-Guevara and Araya-Salas, 2014). Even from the earliest observations, curvature has been synonymous with specialization; we expect curvature to limit the range of functional taxa in a plant-pollinator mutualism and strengthen interactions between the existing participants.

More recently the scope of plant-hummingbird research has expanded to address the biogeography of curvature. As predicted by Stiles (2004), Maglianesi (2015a) and Sonne (2019) found plant-hummingbird curvature to be more represented across species in the lowlands of the Neotropics compared to higher elevations. In this case, plant-pollinator curvature is a form of niche divergence evolving in species-rich lowland habitats, where species experience relatively higher competition (e.g. for nectar or pollen vectors) than in the adjacent Andes mountains (Stiles, 2004; Graham et al., 2009). Furthermore, because plant and hummingbird morphology is better matched (i.e. more specialized) at lower latitudes (Sonne et al., 2020) and hummingbirds with curved bills are predominately tropical, we might expect the occurence of curvature in these taxa to have a predictable latitudinal distribution.

While curvature mediates specialization in plant-hummingbird systems, in other nectivorous bird groups curvature appears to evolve in the absence of selection for resource partioning. In plant-passerine systems, curvature is more prevalent in pollinators than in flowers. Straight flowers do not necessarily exclude pollination by curve-billed birds; for example, the straight, tubular flowers of African *Aloe* L. (Asphodelaceae) are pollinated by curve-billed sunbirds (Paton and Collins, 1989), and the small campanulate flowers of *Vaccinium* L. (Ericaceae) are pollinated by the Hawaiian honeycreeper *Drepanis coccinea* (Fringillidae, Carothers, 1982). For these passerine clades, a dietary shift to nectivory may drive the evolution of bill curvature because (ancestral) insect-pollinated plants require perching at angles not directly facing the flower opening (Paton and Collins, 1989). Furthermore, probing concealed nectar from a fixed perch is an inherently arc-like motion (analogous to reaching into a tall cup to extract, say, an ice cube). Recent experimental work found that Amethyst sunbirds (*Chalcomitra amethystina*) extract nectar more efficiently when flowers are curved towards a perch, indicating that bill curvature may evolve from the inherent arc-like motion of probing tubular flowers (Johnson et al., 2020).

A comparison of honeyeaters (Meliphagidae; predominately Australasian), sunbirds (Africa, Australasia), Hawaiian honeycreepers, and hummingbirds (Nearctic, Neotropic) suggests that bill curvature is widespread in the passerine families but is generally restricted to a subfamily of hummingbirds (Phaethornithinae, Paton and Collins, 1989). While hummingbirds possess unique musculature and wing shape for hovering and manoeuvring adeptly while feeding (Dakin et al., 2018), passerines typically perch to probe for nectar. For plant-passerine systems, primarily insectivorous birds may have evolved curved bills to feed at awkward angles from plants with straight, tubular flowers, and only in some cases does reciprocal adaptation produce curved flowers, e.g. sunbird-pollinated *Streptocarpus dunnii* Mast. (Gesneriaceae, Hughes et al., 2007). The evolution of reciprocal curvature in such plants may be driven by selection to either exclude inefficient pollinators such as bees, which otherwise would compete for nectar and pollen, or to increase the precision of pollen placement. It is worth noting that in landbirds, the evolution of bill shape is coupled with skull shape (Bright et al., 2016; Navalón et al., 2020). Therefore, for some nectivorous passerines the evolution of bill shape is influenced not only by flower morphology, but also the biomechanical factors that restrict skull shape (Navalón et al., 2020). Thus, unlike plant-hummingbird systems (in which curvature mediates pollinator partitioning), selection for curvature in plant-passerine systems appears to operate on nectivorous birds moreso than the plants they pollinate.

Floral diversity contributes to floral isolation and diversification in the angiosperms (Kay and Sargent, 2009; Armbruster and Muchhala, 2009; Vamosi et al., 2018). Similarly, dietary specialization within pollinator clades has contributed to the diversification of mouthpart morphology (Weinstein and Graham, 2017; Maruyama et al., 2018). In both cases, curvature is a widespread feature of morphological diversity. Therefore, to synthesize our knowledge of curved plant-pollinator systems, curvature is a concept that needs an exact definition and method of measurement. In the following section we summarize the approaches used in measuring curvature within the field of pollination ecology, and evaluate the strengths and shortcomings of each. Building on this assessment, we offer a conceptualization of curvature that improves the precision of measurement of this trait. Although this review is motivated by the problem of measuring curvature in plant-pollinator systems, the solution is general to any biological form modelled as a line curve: we then apply this method to floral curvature in an example demonstration.

## Summary of the literature: history of measuring curvature in pollination ecology

We searched the scientific literature for studies of floral or pollinator mouthpart curvature, as these traits are commonly measured as a proxy for specialization. We make the distinction between measuring curvature in a single plane (e.g. the dorsiventral plane of flowers) versus the curvature of surfaces. While single plane images are analysed for line-curvature, measurements of specimens measured in two planes (e.g. dorsiventral and transverse) can be used to analyse surface (Gaussian) curvature (Nath et al., 2003; Coen and Rebocho, 2016). At present, surface curvature has yet to be considered in the context of pollination, and is probably of limited importance. Floral tubes are generally circular in cross-section and surface curvature takes the form of simple or flared cylinders, and complex and mathematically interesting curved surfaces such as saddle-shapes, are absent. Furthermore, because line and surface curvature are related mathematical concepts, it will benefit pollination research to clarify the simplest case (lines), with the goal of generating interest in related ideas, including the curvature of surfaces.

The literature was sourced by querying Web of Science and Google Scholar for a topic search of (curv*) AND (pollinat*) AND (flower OR corolla OR *bird OR *bee OR moth OR *fly). The initial search returned over 300 studies that were then screened for those that measured curvature of floral organs (e.g. petals, styles) and/or animal mouthparts (e.g. bird bills, moth tongues). We sorted studies based on the criteria that 1) the study focused on petal curvature or animal pollination, including qualitative measures of curvature or 2) the study measured curvature of a floral organ other than petals (e.g. style curvature in autogamous species) or animal mouthparts outside of a pollination context (e.g. taxonomic classification).

Under the first criterion, 45 studies were identified to have used some form of curvature metric (Table 1). An additional 17 studies (second criterion) are included in Table S1. There were numerous studies of plant-animal morphology that did not address curvature — these were omitted from our analysis.

**Table 1:** Summary of literature reviewed for metrics of floral or mouthpart curvature in plant-pollinator systems.

| Citation | System | Stated or Inferred<br>Method |
| --- | --- | --- |
| Snow and Snow<br>(1972) | Bill morphology and niche partitioning of nine hummingbird species in the Arima Valley, Trinidad. | qualitative |
| Stiles (1975) | Corolla morphology of *Heliconia* (Zingerberales) and bill morphology of nine hummingbird species at La Selva, Costa Rica. | qualitative |
| Feinsinger and<br>Colwell (1978) | Bill morphology and niche partitioning within hummingbird communities of the Carribean Islands and Monteverde, Costa Rica. | qualitative |
| Gill and Wolf<br>(1978) | Sunbird bill diversity and abilities to extract nectar from Kenyan *Leonotis nepetifolia* (Lamiaceae). | mandibular index |
| Carothers (1982) | Variation in bill morphology in three species of Hawaiian honeycreeper, and effects on feeding performance from *Vaccinium calycinum* (Ericaceae). | angle of deflection |
| Grant and Grant<br>(1983) | Effects of Hawkmoth proboscis length on<br>pollination of *Mirabilis longiflora*<br>(Nyctaginaceae). | qualitative |
| Mountainspring<br>(1987) | Sexual dimorphism and foraging preferences of<br>the Maui Parrotbill (*Pseudonestor xanthophrys*). | mandibular index |
| Paton and Collins<br>(1989) | Functional ecology of bill shape in<br>hummingbirds, honeyeaters, sunbirds, and<br>Hawaiian honeycreepers. | mandibular index |
| Muller (1995) | Curved bristles on the proboscis of European bees<br>for the extraction of pollen. | qualitative |
| Smith et al. (1995) | Correlated evolution of diet and bill shape in<br>Hawaiian honeycreepers. | inverse radius |
| Stiles (1995) | Effects of bill morphology on insect foraging<br>strategy by 11 species of hummingbirds at La<br>Selva, Costa Rica. | arc:chord ratio |
| McIntyre and<br>Browne (1996) | Phototropism in *Helianthus* (Asteraceae) and<br>effects on cotyledon curvature. | angle of deflection |
| Manning and<br>Goldblatt (1997) | Comparative floral morphology of fly-pollinated<br>Iridaceae, Geraniaceae, and Orchidaceae of South<br>Africa. | qualitative |
| Cotton (1998) | Survey and description of 16 hummingbird species occurring at Amacayacu National Park, Colombia. | qualitative |
| Temeles et al. (2000) | Sexual dimorphism of bill shape in Purple-throated caribs (*Eulampis jugularis*) and effects on pollination of *Heliconia* at Quilesse Reserve, Saint Lucia. | angle of deflection,<br>inverse radius |
| Borgella Jr et al. (2001) | Effects of bill morphology (21 hummingbird spp.) on pollen loads (35 plant spp.) at Coto Brus, Costa Rica. | arc:chord ratio |
| Temeles and Kress (2003) | Sexual dimorphism of bill shape in Purple-throated caribs (*Eulampis jugularis*) and effects on pollination of *Heliconia* at Saint Lucia and Dominica. | angle of deflection |
| Travers et al. (2003) | Nectar spurs of *Impatiens* spp. and Ruby-throated hummingbird (*Archilochus colubris*) at Franklin County, Massachusetts. | angle of deflection |
| Temeles et al. (2005) | Sexual dimorphism of bill shape in Purple-throated caribs (*Eulampis jugularis*) and effects on pollination of *Heliconia* at Quilesse Reserve, Saint Lucia. | angle of deflection,<br>inverse radius |
| Collins (2008) | Foraging efficiency from artificial and natural (15 spp.) flowers by 4 species of hummingbirds at Monteverde, Costa Rica. | mandibular index |
| Stiles (2008) | Correlations of bill morphology to the elevational distributions of 150 spp. of hummingbirds in the Andes. | arc:chord ratio |
| Young (2008) | Effects of spur shape on male and female fitness in *Impatiens capensis* (Balsaminaceae). | angle of deflection |
| Martén-Rodríguez et al. (2009) | Testing the pollination syndrome hypothesis in Antillean Gesneriaceae. | angle of deflection |
| Temeles et al. (2009) | Effects of natural (*Heliconia*) and artificial flower morphologies on foraging performance of Purple-throated caribs (*Eulampis jugularis*) at Saint Lucia. | angle of deflection,<br>inverse radius |
| Luo and Li (2010) | Effects of light and temperature on style curvature in *Alpinia* (Zingiberaceae). | angle of deflection |
| Temeles et al. (2010) | Sexual dimorphism of bill shape in 21 species of Central and South American hummingbirds. | angle of deflection,<br>inverse radius |
| Berns and Adams (2010) | Sexual dimorphism of bill shape in black-chinned hummingbird (*Archilochus alexandri*) and ruby-throated hummingbird (*Archilochus colubris*). | geometric<br>morphometrics |
| Berns and Adams<br>(2013) | Sexual dimorphism of bill shape in 219 hummingbird spp. | geometric morphometrics |
| Wang et al. (2013) | Selection for nectar spur curvature in *Impatiens oxyanthera* (Balsaminaceae) mediated by pollinators and nectar robbers. | angle of deflection |
| Maglianesi et al.<br>(2014) | Elevational variation in plant-hummingbird network structure mediated by bill morphology. La Selva, Costa Rica. | angle of deflection |
| Rico-Guevara and<br>Araya-Salas (2014) | Selection for bill shape driven by male-male competition in *Phaethornis longirostris*. La Selva, Costa Rica. | arc:chord ratio |
| Alexandre et al.<br>(2015) | QTL analysis comparing hummingbird pollinated and generalist *Rhytidophyllum* flowers (Gesneriaceae). | angle of deflection |
| Campos et al. (2015) | Generating 3D printed flowers to experimentally study the feeding mechanics of moth pollination. | curve decay parameter |
| Maglianesi et al.<br>(2015b) | Differential preferences of artificial and natural (65 spp.) flower populations visited by 3 species of hummingbird in Braulio Carrillo National Park, Costa Rica. | angle of deflection |
| Maglianesi et al. (2015a) | Plant-pollinator specialization along an elevational gradient at Braulio Carrillo National Park, Costa Rica. 21 hummingbird spp. and 208 plant species examined. | angle of deflection |
| Rocha et al. (2015) | Physical and biochemical basis for androgynophore bending in bat-pollinated *Passiflora mucronata* (Passifloraceae). | arc:chord ratio |
| Miller et al. (2017) | Ecological divergence among closely related, morphologically similar Honeyeaters co-occurring in arid Australian environments. | arc:chord ratio |
| Lagomarsino et al. (2017) | Evolution of pollination syndromes in Andean Campanulaceae. | arc:chord ratio |
| Boehm (2018) | Review of nectar robbing in *Centropogon* (Campanulaceae) | qualitative |
| Hadley et al. (2018) | Effects of forest fragmentation on hummingbird bill morphologies (19 spp.) representative of specialization. Coto Brus, Costa Rica. | arc:chord ratio |
| Joly et al. (2018) | Testing the pollination syndrome hypothesis in Antillean Gesneriaceae. | geometric<br>morphometrics |
| Partida-Lara et al.<br>(2018) | Spatio-temporal structure of the taxonomic and functional diversity of hummingbirds at the biosphere reserve El Triunfo, Chiapas, Mexico | inverse radius |
| Dellinger et al.<br>(2019) | Floral trait changes correlated with the repeated shifts away from buzz pollination in the Melastomataceae. | qualitative |
| Peng et al. (2019) | Evaluation of fitness optima of the moth proboscis and flower shape. | curve decay<br>parameter |
| Sonne et al. (2019) | Variation strength of plant-hummingbird specialization along an elevational gradient in Podocarpus National Park, Ecuador. | arc:chord ratio |
| Johnson et al. (2020) | Experimental effects of floral orientation and curvature on nectar extraction by Amethyst sunbirds (*Chalcomitra amethystina*). | angle of deflection |

In our survey, the dedicated discussion of dorsiventral curvature in plant-pollinator interactions begins with Hainsworth (1973), in reference to *Heliconia* and hermit hummingbirds. Curvature in pollination ecology is first empirically studied by Gill and Wolf (1978), though methods for measuring curvature of bird appendages outside of a pollination context can be found much earlier (Baldwin et al., 1931). We identified six common approaches to measuring curvature in pollination systems. These are:

1. *qualitative description*, e.g. “very curved,” “less curved,” but these are generally no longer used;
2. the *arc:chord* method defines curvature as a ratio of two lines: an arc fitted to the curve of a flower or mouthpart (e.g. bird bill) from its tip to base, and the straight line (chord) subtending the arc (Figure 1);
3. the *mandibular index* defines curvature as a ratio of two lines: a straight line from base to tip (chord) and a perpendicular line that measures the maximum height of the flower/bill arc (versine).
4. the *angle of deflection* method defines curvature as the angle between the tangent line at the base of the flower/bill and the straight line from base to tip (chord);
5. the *inverse radius* method which approximates the arc of the flower/bill as a segment of a circle. Curvature is defined as the inverse radius of the fitted circle;
6. *geometric morphometrics*, which quantifies shape as a configuration of homologous points (landmarks) existing on a coordinate plane (Figure S2).

**Figure 1.**
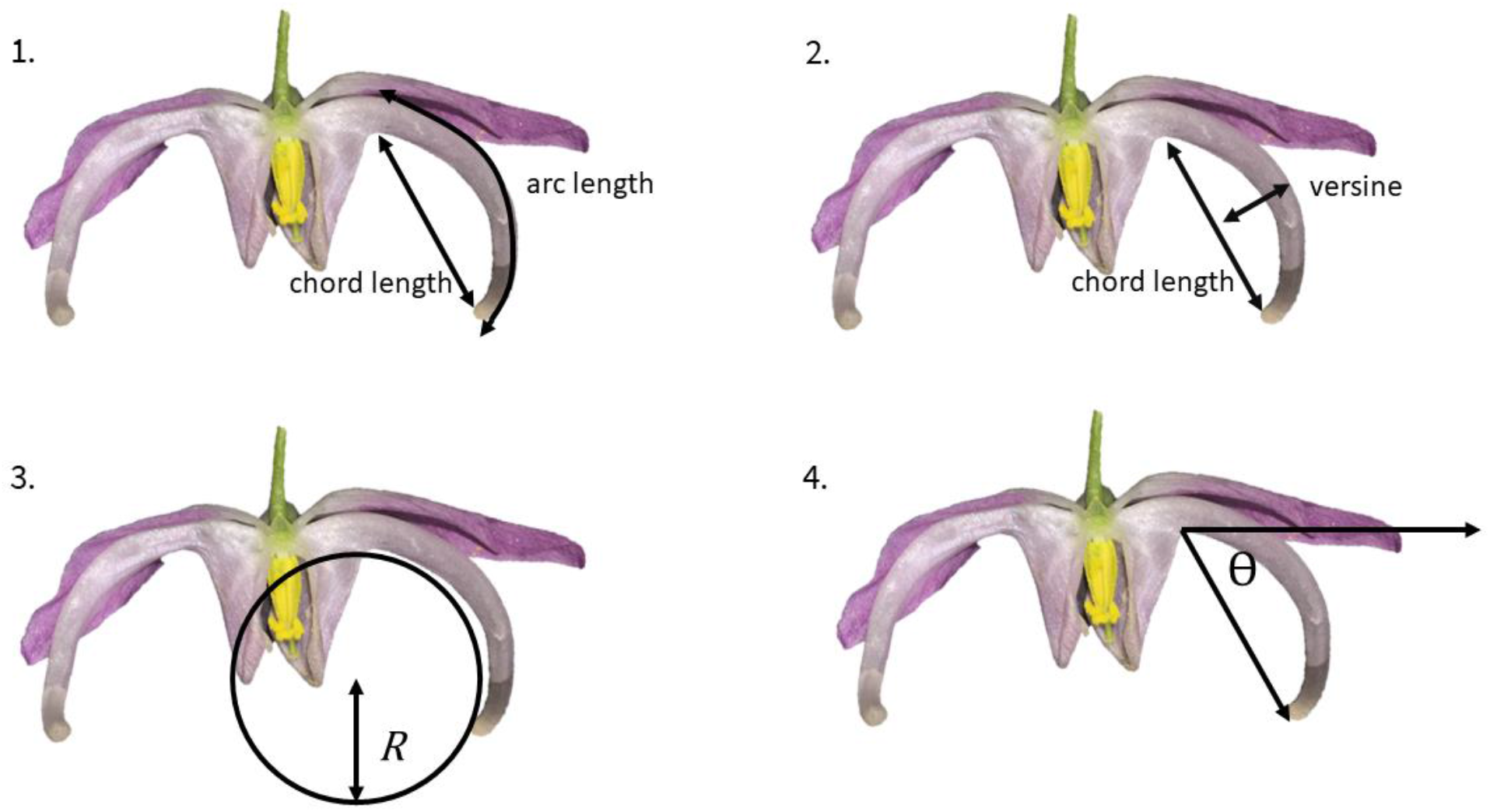
Overview of most commonly used curvature metrics within pollination ecology. 1.arc:chord ratio. 2. mandibular index 3. inverse radius. 4. angle of deflection

The strength of methods 2–5 are their portability and accessibility. These measurements can be taken in the field or determined easily from photographs. The methods are intuitive and in the simplest case, require only a ruler, string, and protractor. Temeles (2009) pointed out that, for curves well fitted by a circle, the *inverse radius* method is interchangeable with the *angle of deflection* method because the radius can be calculated from the length and angle of a chord (tangent chord theorem, Bell, 1956). Similarly, for circles, arc length and versine can be computed from the radius and angle of deflection (Zwillinger, 2018). Given the required additional geometric information, for curves well fitted by a circular arc, methods 2–4 will be proportional to the *inverse radius*.

In fitting a curve with a circle we assume that curvature is constant across the specimen. However, when curvature deviates from constant, methods 2–5 are less suitable (discussed in Berns and Adams, 2010). The *angle of deflection* and *mandibular index* are computed from only two or three landmarks, respectively. Any changes in curvature between landmarks are not considered—this is problematic when curvature varies across the specimen (e.g. a hook at the end of a bird bill). The *inverse radius* method assumes the curve to have constant curvature. The *arc:chord* method, because it considers arc length, implicitly contains information about curvature across the entire specimen—for a curved line in a plane, arc length and total curvature are proportional (Section 3). However, the curvature of a local feature cannot be extracted given only the length of the arc and chord. Overall, while these methods succeed in estimating the total curvature of an entire curve, they are not designed to account for fluctuating curvature caused by locally curved features.

An additional problem is a lack of consensus terminology and methodology. For example, the *arc:chord* method is also called the *maxillary index*, while the *angle of deflection* method is sometimes referred to as the *angle of declension* method. Many studies create their own terminology for the concept of arc length: the length of a curve between two points. Most studies define their own terms for measuring and reporting curvature without reference to previous studies. We found no discussion of units or of their meaning. This creates uncertainty about how to compare and convert metrics used between studies. We also found no discussion or methods accounting for allometry. We believe these problems could be remedied by referring to the mathematical literature for the derivation and definition of curvature and related concepts (see: Section 3).

Starting with Berns and Adams (2010) geometric morphometrics (GM) emerges in the pollination literature. GM is broadly concerned with the covariance of shape and associated or causal variables (Bookstein, 1991; MacLeod, 2002), for example, how flower shape might covary with local pollinator communities (Gómez et al., 2009). This approach has steadily gained in popularity due to its mathematical rigour, reproducibility, and the appealing visual representations of shape variation (Olsen, 2017). Additionally, because GM has a traceable mathematical lineage (Bookstein, 1991), its vernacular is well-defined and used consistently between practitioners. Here, we highlight some of the most important features of GM to introduce relevant concepts, and recommend the concise and authoritative introduction by Webster and Sheets (2010) for more details.

In *traditional* morphometrics univariate measurements (such as length, width, angle) are the primary data used to quantify shape. Methods 2-5 (above) fit into this category. Typically these measurements are anchored by landmarks—toplogically or biologically homologous points that can be located on all specimens (detailed in Bookstein, 1991; MacLeod, 1999). In *geometric* morphometrics landmarks are assigned across the specimen, with the goal of representing its shape as completely as possible. A typical protocol for a 2-D object begins by placing the specimens on an *xy*-grid and assigning *xy*-coordinates to landmarks (Figure S2). In a comparative study, the samples are overlaid so that their shape information is isolated from their orientation, location, and size. This is done using a least-squares type protocol, most commonly the Generalized Procrustes Analysis (Rohlf and Slice, 1990). In *outline-based* GM, the *xy* coordinates of landmarks are fitted by form or shape functions (sensu MacLeod, 2012) and decomposed by elliptic (Kuhl and Giardina, 1982) or ‘ZR’ (Zahn and Roskies, 1972) Fourier analysis, respectively. Harmonic shape variables from a Fourier analysis are then used to calculate the principal components of shape variation (MacLeod, 2012). In *landmark-based* GM, the set of landmarks summarising the shape of an organism is treated as a ‘landmark configuration.’ Configurations exist in a shape space defined by the number of landmarks and spatial dimensions implemented. These configurations are then projected onto a simpler Euclidian space, analogous to the reduction of a spherical Earth onto a two-dimensional map (Webster and Sheets, 2010). From here, familiar statistical procedures (e.g. PCA) can be performed to quantify variation in landmark configurations (shape) between samples.

A typical presentation of a shape-PCA attaches end-member specimens (MacLeod, 2002) or end-member deformation grids (Bookstein, 1991) at both ends of a given principal axis. This enables a qualitative description of the primary trait(s) varying along said axis. For example, visual inspection of deformation grids along PC1 of Figure 4 (Section 4) could be interpreted as shape variation driven by differences in floral curvature. However, the limitation of *landmark-based* GM in the quantification of curvature is that this method is concerned with analyzing the entirety, not segments, of a specimen’s shape. *Outline-based* GM, though well-suited to analysing open curve segments (MacLeod, 1999), lacks a method for specifically extracting information about curvature (detailed in Section 4). Therefore, while GM has enabled the quantification of shape (sensu Bookstein, 1991), we are currently limited to describing curvature by visual inspection of the principal axes of shape space.

## Curvature: concepts from differential geometry

Reviewing the literature leads us to ask, “what is curvature?” In related fields, for instance in plant physiology, there have been uses of a point-wise definition of curvature resembling that used in differential geometry (Castle, 1962). However, as in pollination ecology, references to the mathematical literature are missing. Therefore, we propose turning to the field of geometry in order to develop the concept of curvature starting from first principles. There, we again find several definitions resulting from a history of independent derivations (reviewed in Coolidge, 1952; Bardini and Gianella, 2016). Nonetheless these definitions share a conceptual theme; curvature is a local property that can be measured point-wise on a line. This concept is fundamentally different from those typically used in pollination ecology, where curvature is a single property of an entire shape. Here we follow the conventions of Casey (1996) and Rutter (2000) and present a definition of curvature that is tractable for analyzing biological shapes.

Intuitively, when a line deviates from being straight we say it is curved, the extent to which it is not straight is its curvature. More technically, a line deviates from being straight when its slope (i.e. the graph of the first derivative) changes magnitude—this is represented here by the rotating unit tangent vectors **T**_0_, **T**_2_, **T**_7_, **T**_*n*_ in Figure 2. Therefore, curvature can be thought of as the rate of change in the tangent as we move across the curve. Hence, the tangents of a straight line will have the same slope everywhere, and the line will have a curvature of zero, whereas when the slopes of the tangents of a curve (Figure 2) fluctuate, the line will have non-zero curvature.

**Figure 2.**
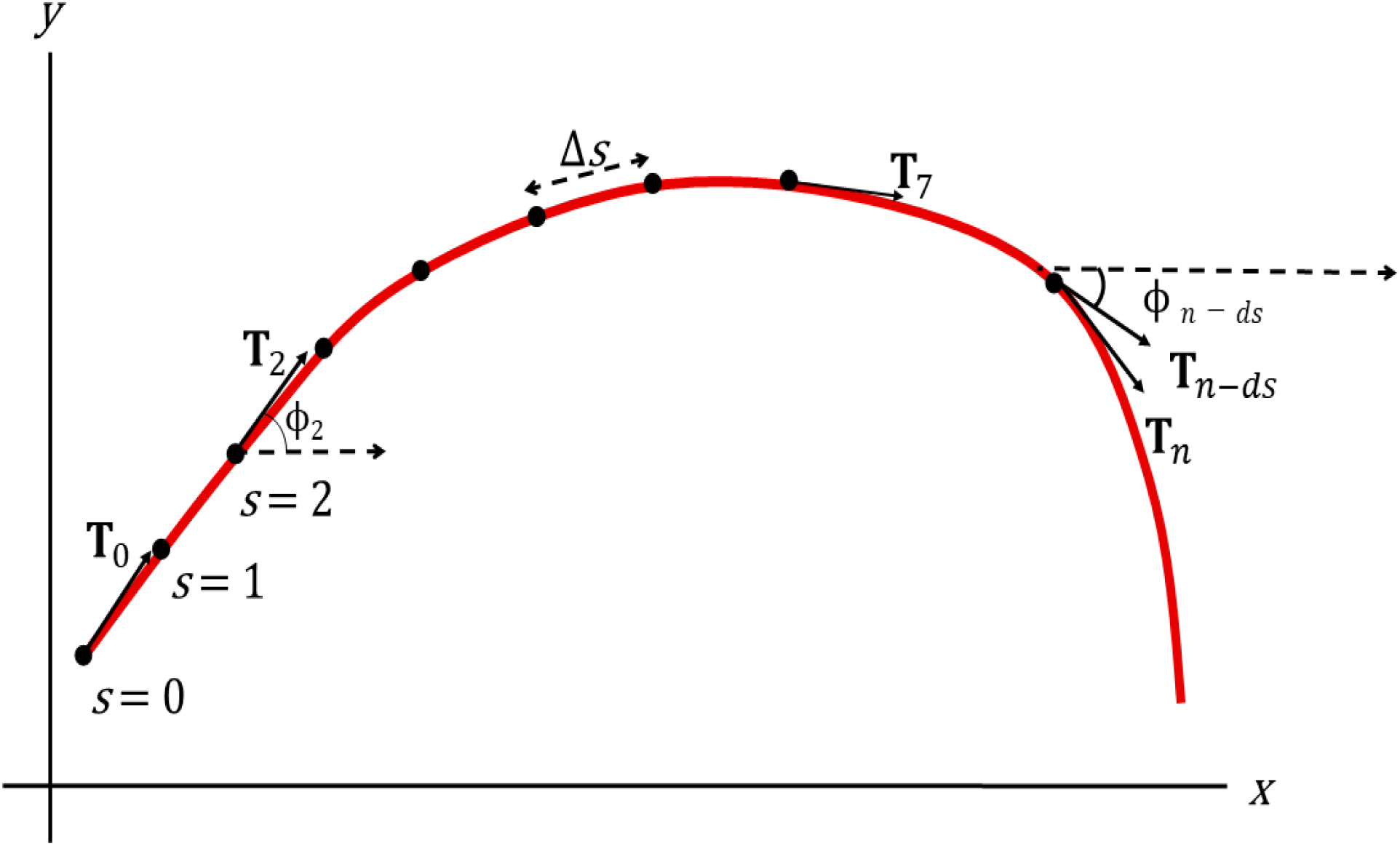
A curve parameterized by arc length, *s*. ***T***_0_, ***T***_2_, and ***T***_7_ are the tangents 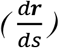 at *s* = 0, *s* = 2, and *s* = 7, respectively. For convenience the tangents are displayed as indications only, not exact representations. *Φ*_2_ and *Φ*_*n*–*ds*_ represent the angles formed between the tangent **T**_2_ and the *x* axis, and *T*_*n*–*ds*_ and the *x* axis, respectively. Total curvature is the sum of the changes in rotation (*ΔΦ*) along the curve.

As biological curves often loop back on themselves (e.g. spirals) they are best described by parametric functions. By using a ‘hidden’ variable that determines the values of *x* and *y* independently, parametric functions allow a curve to have multiple *y* values for a single *x*. Here, we use the parameter variable arc length, *s*, along the curve, to give us the *x* and *y* position. Specifically, we can express a position vector **r** = [*x,y*] as a function solely of arc length, *s*. Using vector notation we have:

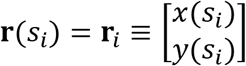

Here **r**_*i*_ is shorthand for **r**(s_*i*_), which indicates that our position (*x*(*s_i_*),*y*(*s_i_*)) on the curve is determined by the length of the segment *s_i_*. Although we could parameterize a curve by many potential parametric variables, arc length is a convenient choice because it allows us to move along the curve at uniform increments, which we denote as *Δs*. This proves useful when taking repeated, equally-spaced measurements (such as curvature) along a curve.

As we are interested in the derivative properties of our arc-length parameterized curve, we can differentiate **r**(*s*) with respect to arc length *s* in the following way (using the formal definition of the derivative):

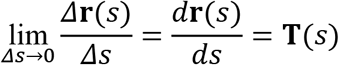

This produces a tangent function 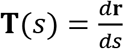 giving the first derivative of the parametric equation **r**(*s*). The tangent **T**(*s_i_*), represented by the shorthand **T***_i_*, contains information about the direction of the curve at position **r***_i_* that we can use to calculate curvature. When the tangent **T** is placed into a cartesian plane, we can re-parameterize by the angle *Φ* formed with the *x*-axis (Figure 2). Thus the *x*’(*s_i_*) and y’(*s_i_*) components of the tangent vector **T_i_** can be expressed as:

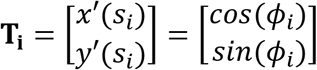

Where:

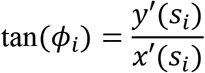

And:

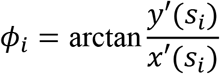

At the beginning of this section we defined curvature, *κ*, as the rate at which the tangent is changing direction. Thus, curvature *κ* can be expressed as the change in the angle *Φ* formed between the tangent **T** and the *x*-axis:

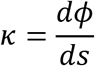

This definition provides an intuitive unit of measurement for reporting curvature: degrees of rotation per unit arc length (Figure 2). For example, if curvature has been calculated at every millimeter along the length of an arc, we would report its curvature as degrees per millimeter *degrees* · *mm*^−1^. Framed this way curvature is a measure of rotation per unit distance. In contrast to previous definitions, where curvature is an indivisible, single property of an entire shape, here, curvature is a property of every measured point along the curve. Under this point-wise definition, we can summarize the *total curvature* (Milnor, 1954) of a specimen as the total point-wise individual curvature along the curve:

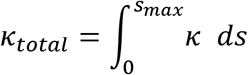

Units for *total curvature* are no longer expressed as *degrees* · *mm*^−1^ because we are not measuring curvature at a single point. Instead we are integrating all tangent rotations along the curve, expressed simply as *degrees*. For two curves of the same shape (*sensu* Bookstein, 1991), their total curvature will be equal, regardless of allometric scaling. For example, compare the half-unit circle with another half-circle with *r* = 2. Although the second curve is larger, the total curvature of both is *π radians*. For comparative studies that need to account for size, we suggest dividing total curvature by arc length, *s*, so that *adjusted total curvature* is:

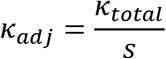

Using the example above, the half-unit circle would have 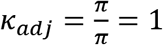 radians, while the half-circle with *r* = 2 would have 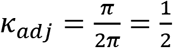 *radians*. Shape being equal, smaller curves will have greater adjusted total curvature. However, the use of this adjustment depends greatly on the biological context of the research question (Klingenberg, 2016).

This concept of curvature has been embedded within the morphometrics literature since the 1970’s. The widely-applied tangent angle function (*Φ*(*t*)) defined by Zahn and Roskies (1972) describes complex shapes by measuring the tangent angle (*Φ*) many times along a shape’s perimeter. This approach to describing shape is analagous to following cardinal directions in wayfinding. Starting from the first landmark, the tangent angle (*Φ*) between landmarks is calculated step-wise around the shape. This step-wise progression eventually outlines the shape’s perimeter (MacLeod, 2011). The resultant tangent angle graph is a unique single-valued function describing the specimen’s shape and implicitly contains information about point-wise curvature. However, because this technique was developed for quantifying *shape*, its utility as a curvature metric has generally not been recognized or applied (but see: Van Otterloo, 1991). This point is discussed further in the following section.

## A proposed protocol for measuring curvature

As illustrated in our methodology review, the current protocols for measuring flower-pollinator curvature lack conceptual unity. There are two main advantages of the curvature definition described above. First, curvature becomes a local property of the tissue or organ under study. This means that shape information is gathered at every point along the curve and can be examined and compared to other points within or between specimens. This differs from previous methods that take curvature as an indivisible property of the entire specimen. Second, because the revised definition comes directly from the field of differential geometry, we benefit from established, well-defined concepts that make clear what is meant by ‘curvature.’ When the definition of curvature is concordant between these research areas, future advances in geometry can be more readily incorporated into morphological studies.

In order to apply the point-wise definition of curvature, a biological organ or tissue needs to be reduced to a continuous function. To do this, we propose a protocol as illustrated in Figure 3. First, a specimen is landmarked at several locations along the region of study. Second, a mathematical function is fitted to the landmarks, and finally, curvature is calculated point-wise along the curve. The long-term goal of this proposal is to integrate the analysis of curvature with existing morphometrics protocols. In the following demonstration we use existing morphometric tools for landmarking and curve-fitting—these were previously developed in the field of traditional and outline morphometrics (e.g. Rohlf, 1990; MacLeod and Rose, 1993; Terral et al., 2004, Section 2).

**Figure 3:**
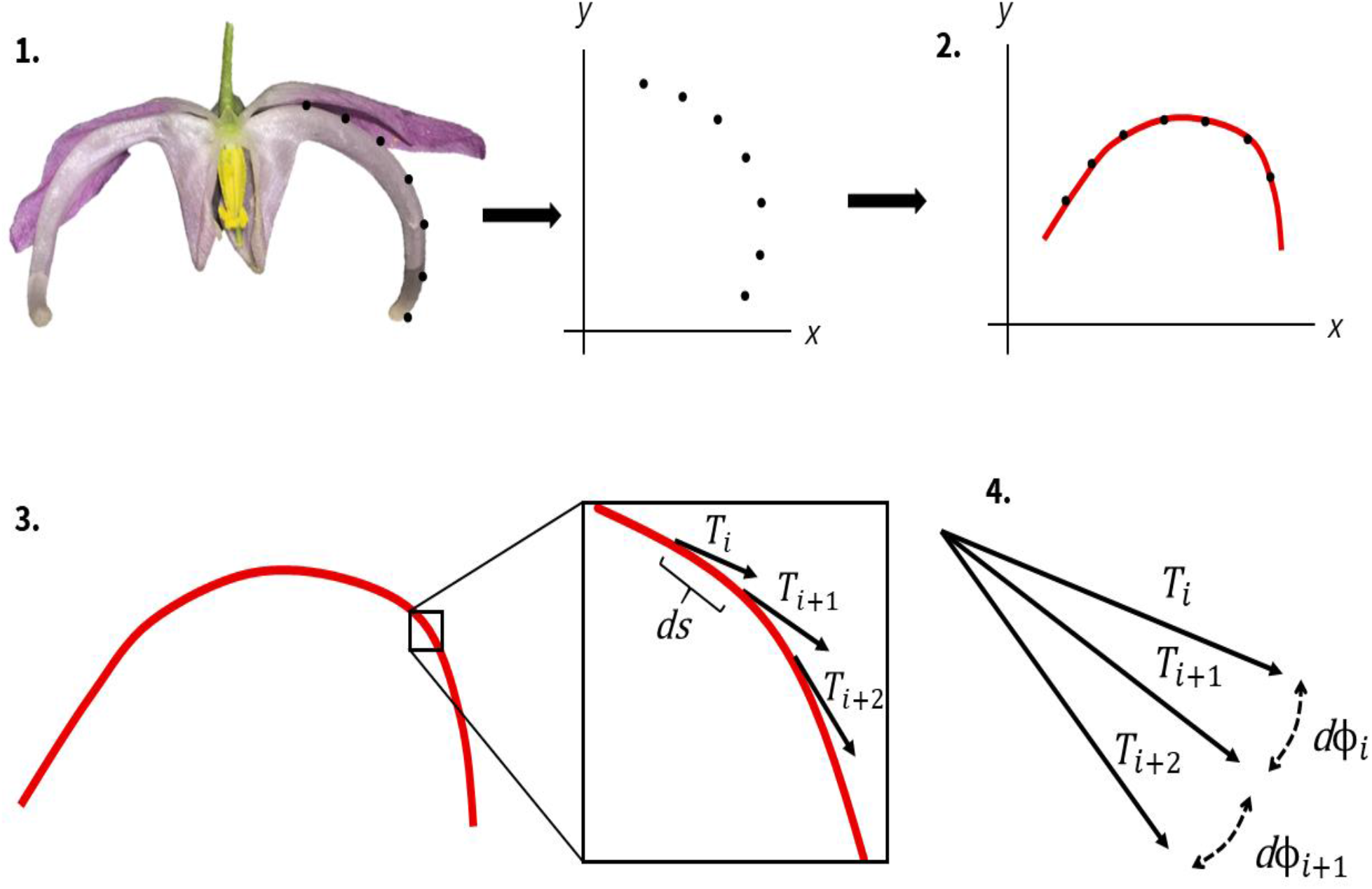
Proposed protocol for measuring curvature. 1. A specimen (in this case Epimedium violaceum) is landmarked, semi-landmarked and assigned xy coordinates within a Cartesian plane. 2. The xy coordinates are rotated so that a single-valued function can be fitted to the landmarks (this step is not necessary for all cases). 3. The tangent vector **T** is calculated at an arbitrarily large number of increments, ds, along the curve. 4. Curvature is calculated as the rate of change of the tangent angle (Φ) point-wise along the curve. Total curvature is calculated by the methods outlined in Section 3.

As mentioned in the previous section, the field of plant physiology was an early adopter of the point-wise definition of curvature. In one case we found curvature (as defined above) computed from cubic functions fitted to cucumber seedlings that had been landmarked by hand (Cosgrove, 1990). Our protocol can be seen as a computerized version of this procedure. In another study, total curvature of *Anthurium* Schott (Araceae) spadices was computed from fitted B-spline curves (Pour et al., 2018). However, because landmarks were not defined and the scripts are not publicly available, the reproducibility of this protocol is low. Here, we propose to develop the analysis of curvature specifically within the R programming environment (R Core Team, 2017), where existing landmarking and curve-fitting procedures can be called, and where modern morphometrics is being most actively developed (e.g. Adams and Otárola-Castillo, 2013; Bonhomme et al., 2014).

In the following section we calculate curvature from cubic smoothing splines (Wood, 2017) fitted to landmarks. While splines are suitable for relatively simple, open curves, we encourage the development of algorithms that will compute curvature from more sophisticated curve-fitting strategies (reviewed in Rohlf, 1990; MacLeod, 2002). Already there is potential for existing GM algorithms to be modified to compute curvature. Notably, the tangent angle function, *Φ*(*t*), describes the angle of the tangent at a landmark as a function of distance traveled along the specimen’s outline (Zahn and Roskies, 1972; MacLeod, 2012). While the concept of *Φ* (*sensu* Zahn and Roskies, 1972) is equivalent to the one described in Section 3, the tangent angle function appears to have been derived without reference to differential geometry (Raudseps, 1965). Though we are interested specifically in the curvature of shapes, the tangent angle function was developed as a means to quantify shapes in their entirety—curvature is incidental. Nonetheless, when many interpolated semi-landmarks are used to model a specimen’s outline (e.g. Zahn and Roskies, 1972), the tangent angles could be summed to give an approximation of *K_total_*. There is great opportunity for existing tangent angle algorithms (e.g. Claude, 2008) to parse curvature data as a part of an outline morphometrics analysis.

## Demonstration: A study of the development of curvature in *Epimedium*

### Study System and Methods

In this demonstration, we compare the development of floral morphology in two subspecies from the *Epimedium grandiflorum* complex (Stearn, 2002): *Epimedium koreanum* Nakai (*Epimedium grandiflorum* var. *koreanum* (Nakai) K.Suzuki), and *E. violaceum* C.Morren & Decne. (*Epimedium grandiflorum* f. *violaceum* (C.Morren & Decne.) Stearn). Though closely related, these taxa have notable differences in floral pigmentation and size (Stearn, 2002). We investigated whether there were consequent differences in floral *shape*, including curvature of the prominent nectar spurs. This study serves to demonstrate that the analysis of curvature can be improved by clearly defined protocols and units of measurement, in addition to encouraging the integration of traditional and geometric morphometrics.

Following an initial description of *Epimedium* development (Supp. Methods), a set of nectar spurs (*n*=57, Table S2) of varying maturity were sampled for imaging. Spurs were photographed in the dorsiventral view using a stereo microscope at 6.3x. Flower shape was quantified from the images using a standard GM protocol in `geomorph` v.3.2.0 (Adams and Otárola-Castillo, 2013), as outlined in Section 4 (see also: Supp. Methods). Shape data was then analysed for the principal components (PCs) describing the major axes of variation in shape space (Figure 4).

**Figure 4:**
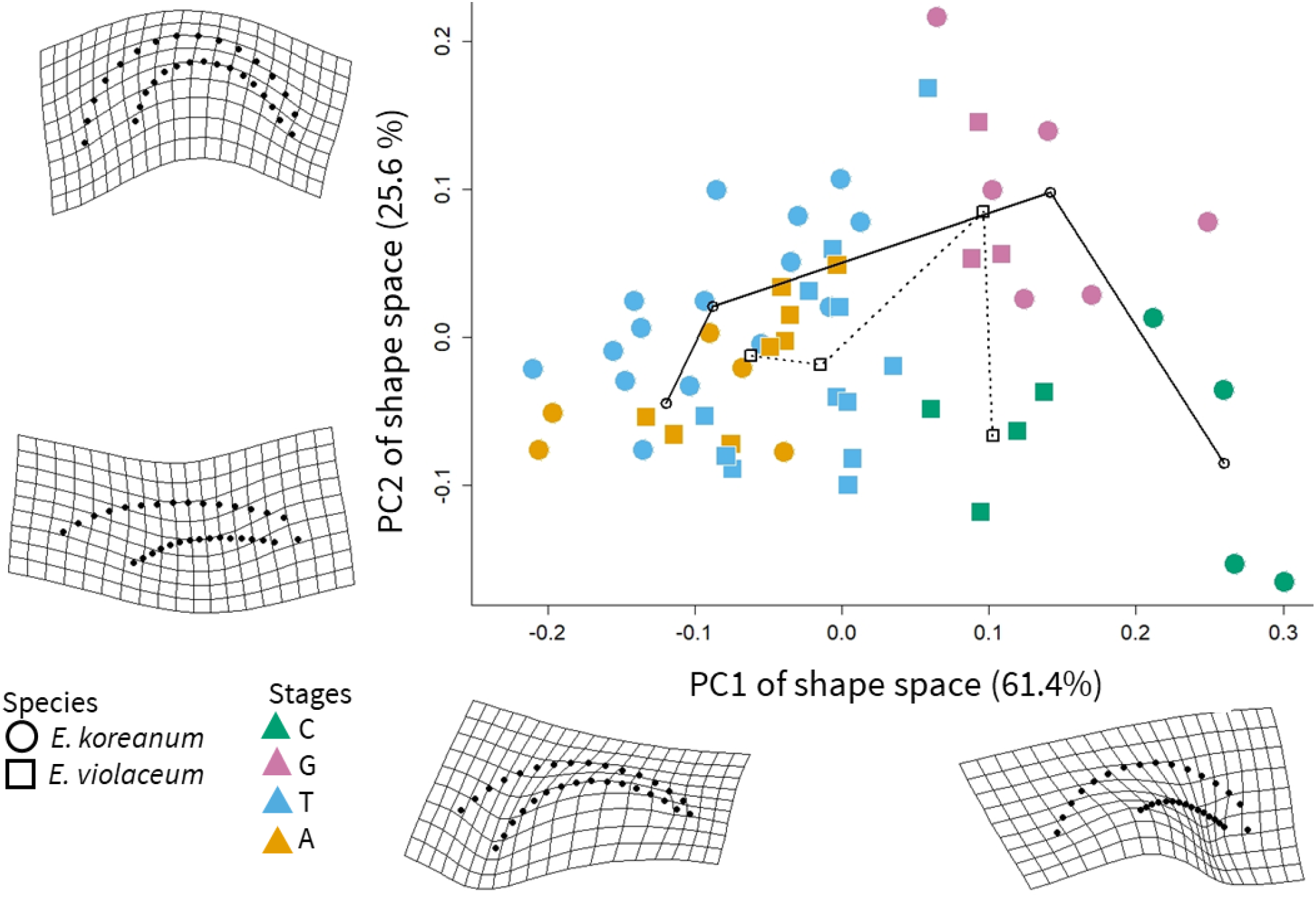
Morphometric trajectory analysis of nectar spur development in E. koreanum (circles) and E. violaceum (squares). The PC axes and deformation grids illustrate the two most explanatory components of shape change in the development of nectar spurs. The two taxa studied occupy significantly different domains of morphospace at the first (green) stage of development. At maturity (yellow), the two species do not differ significantly in shape.

Morphological development can be compared by quantifying the size, orientation, and shape of an individual’s trajectory through morphometric shape space (Adams and Collyer, 2009). We used developmental trajectory analysis (Collyer et al., 2015) to determine how shape varied with growth stage. First, a linear mixed model was fit to relate changes in shape with developmental stage (model variables and likelihood ratio tests for significance are described in the Supp. Materials):

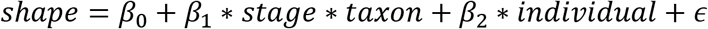

This model was submitted to `geomorph::trajectory.analysis()` to quantify and compare developmental trajectories between the two taxa. Pairwise distances of shape means for each stage were calculated in RRPP v.0.5.2 (Collyer and Adams, 2018).

The deformation grids illustrating variation in PCA shape space suggested that floral curvature increased along both PC1 and PC2 (Figure 4). To analyse curvature specifically, smoothing splines (Cleveland et al., 1992) were fitted to the dorsal side of the nectar spurs using `stats:: smooth. spline()` (Supp. Methods). Total curvature (Section 3) was calculated from the splines using the R package curvr v.0.1.0 (Boehm, 2020). To test for the correlation between the PCs of shape space and total curvature we fit the linear mixed model:

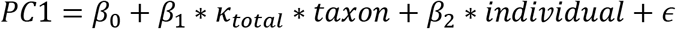

A similar model was fitted to the developmental stage data to test whether total curvature varied with age (Supp. Methods). Estimated marginal means and pairwise comparisons were calculated for each taxon and developmental stage using the `emmeans` package v.1.4.5 (Lenth et al., 2018).

Finally, we re-measured curvature using metrics 2 to 5 as outlined in our summary of the previous literature (Section 2, Supp. Methods), and performed a pairwise regression for each metric, including the point-wise defintion proposed here. Linear measurements (e.g. chord length) were made using tpsDig v.2.3.2 (Rohlf, 2015). The inverse radius was computed from fitted circles using `pracma` v.2.2.9 (Borchers, 2019). In order to estimate deviations from circularity, the root mean square error was computed for each specimen fitted by the inverse radius method.

### Results

Inspection of the deformation grids associated with PC1 and PC2 suggests that curvature is associated with both axes of shape space (Figure 4). Indeed, there is a significant correlation between curvature and PC1 (*p* = 0.019, *t* – *ratio* = 2.40) and PC2 (*p* = 2.1 × 10^−11^, *t* – *ratio* = 8.49, Figure 5). There are no interspecific differences in curvature during floral development, but in *E. koreanum* there is a significant increase in curvature during Stage 2 (Table S6, Figure S6). From mid to late development (Stages 2 to 4) there is 50.6 degrees less total curvature in the nectar spurs of *E. koreanum* (*p* = 0.002, Table S6). Similarly, from early to mid development (Stages 1 to 2) there is 37.9 degrees less total curvature (*p* = 0.073). The specimen photographs (Supp. Materials) suggest that at Stage 2, nectar spurs tend to curve sharply as they elongate (grow) from bud-size spurs (Stage 1) to mature-size. After growth slows, the spurs straighten as they approach anthesis (Stage 4). Whether this developmental pattern is adaptive or incidental is yet unknown.

**Figure 5:**
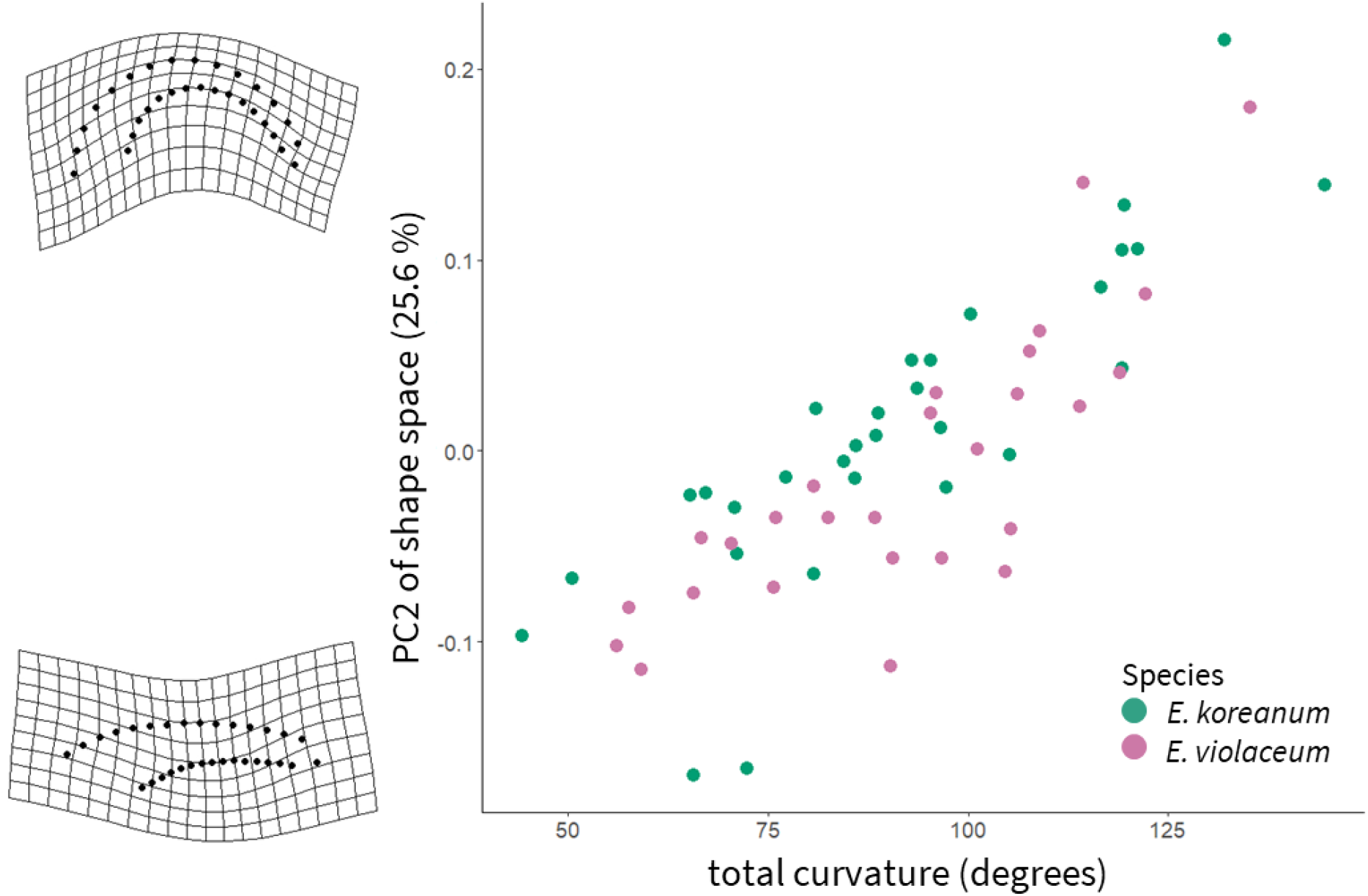
Scatterplot of total curvature and PC2 from PCA of geometric morphometric shape data (see also: Figure 4). PC2 represents the axis of second greatest variation in shape during Epimedium flower development.

Trajectory analysis found that interspecific and temporal differences in shape are driven mainly by PC1, which accounts for 61.4% of shape variation (Table S5, Figure 4). Although both PCs are correlated with curvature, *E. koreanum* at Stage 2 did not necessarily deviate from other samples in morphospace (Table S5). In this case, it may be that other geometric components contribute to spur shape, and curvature alone does not sufficiently explain variation along the PCs of shape space.

Pairwise comparisons of the various curvature metrics found that, for both species, all metrics are significantly correlated (*p* < 0.0043, Tables S8 and S9). The inverse radius had the lowest correlation with the other metrics (Pearson’s *r* < 0.65). Thus, we used `pracma:: circlefit()` to calculate the error distance for each specimen’s fitted circle. Consequently, the specimens approximated poorly by circles had the most residual variation between total curvature and the inverse radius metric (*p* = 3.78 × 10^−7^, *t* = 5.7, Figure S8). This suggests that for non-circular curves, a point-wise metric is better suited to capture local deviations and fluctuations in curvature.

### Discussion and Future Applications

The proposed definition for curvature (as adopted from differential geometry) clarifies both the concept and the units of measurement. Using the terms *point-wise curvature* and *total curvature* enables us to distinguish between the curvature at a point and the cumulative curvature of the structure under study. In the context of pollination ecology, a rigorous measure of curvature will facilitate a number of avenues of investigation. These include matching curvature of flowers and bills in co-evolutionary studies, describing the variation of curvature of flowers and bird bills in populations as raw material for evolution, and investigating the evolution of curved flowers and bird bills in the context of phylogeny.

By pairing an analysis of curvature with geometric morphometrics, we are able to quantify the covariation between curvature and overall shape. Further integration of univariate metrics with GM could be achieved by modifying the principal component analysis of shape to partial out variation due to an explanatory variable (e.g. redundancy analysis, Borcard et al., 2018).

Because GM is ubiquitous among studies of biological form, we hope that this metric for curvature, as presented here, may facilitate communication via a common language between fields that are interested in curved structures. The diversity of applications spans the study of sexual selection on horn, tusk, and antler structure (e.g. in dung beetles Emlen et al., 2005), the twining and nutation of tendrils (Goriely and Neukirch, 2006; Bastien et al., 2014), the functional and behavioural ecology of claw shape (e.g. in birds, lizards, and dinosaurs Birn-Jeffery et al., 2012), the bending of hypocotyls in response to gravity and light (Silk, 1989; Frassen et al., 1982), and the biomechanics of locomotion and escape behaviours (e.g. prehensile tails in Neotropic monkeys, Schmitt et al., 2005; lateral bending of lizards, Ritter, 1992). Widening the applicability of this curvature metric will require drawing deeper from the morphometrics toolkit. Beyond simple curves that might be fitted by polynomials, splines are necessary for describing spirals, undulating lines, and multi-valued functions. For closed outlines, ZR Fourier Analysis (Section 2, Zahn and Roskies, 1972; MacLeod, 2011) is well-suited for decomposing a range of complex biological shapes. As discussed in Section 3, the tangent angle function used to represent a closed outline mirrors the defintion of curvature used in differential geometry. Fortunately, `Momocs` (Bonhomme et al., 2014) offers functions for ZR Fourier Analysis, providing a convienient launch point for integrating point-wise curvature estimation with the quantification of closed outlines. With much of the groundwork already laid, estimating point-wise curvature could soon be applied to a diversity of study systems.

### Conclusions

In this synthesis we discussed the ecological significance of curvature within the field of pollination ecology. In reviewing the methods used to measure curvature, we found a need for both conceptual and methodological unity. By drawing from the geometry literature we aimed to clarify the definition of curvature within the contexts most commonly used in pollination ecology: the curvature of floral tubes or pollinator mouthparts in the dorsiventral plane. We demonstrated the utility of this revised metric by analysing the development of curved nectar spurs in *Epimedium*. For curves that deviate from segments of circles, a point-wise definition of curvature is recommended over historical methods that model circularity on segments of a circle. Furthermore, a clearly defined concept of curvature creates the potential for progressive integratation with the swiftly advancing, and broadly applied field of geometric morphometrics.

## References

Adams, D.C., and Collyer, M. (2009). A general framework for the analysis of phenotypic trajectories in evolutionary studies. Evolution 63, 1143–1154.

Adams, D.C., and Otárola-Castillo, E. (2013). Geomorph: An R package for the collection and analysis of geometric morphometric shape data. Methods in Ecology and Evolution 4, 393–399.

Alexandre, H., Vrignaud, J., Mangin, B., and Joly, S. (2015). Genetic architecture of pollination syndrome transition between hummingbird-specialist and generalist species in the genus *Rhytidophyllum* (Gesneriaceae). PeerJ 3, e1028.

Armbruster, W.S., and Muchhala, N. (2009). Associations between floral specialization and species diversity: Cause, effect, or correlation? Evolutionary Ecology 23, 159–179.

Baldwin, S.P., Oberholser, H.C., and Worley, L.G. (1931). Measurements of birds (Cleveland Museum of Natural History).

Bardini, G., and Gianella, G.M. (2016). A historical walk along the idea of curvature, from Newton to Gauss passing from Euler. International Mathematical Forum 11, 259–278.

Bastien, R., Douady, S., and Moulia, B. (2014). A unifying modeling of plant shoot gravitropism with an explicit account of the effects of growth. Frontiers in Plant Science 5, 136–145.

Bell, J. (1956). Tangent, chord theorem. The Mathematical Gazette 40, 211–212.

Berns, C.M., and Adams, D.C. (2010). Bill shape and sexual shape dimorphism between two species of temperate hummingbirds: Black-Chinned hummingbird (*Archilochus alexandri*) and Ruby-Throated hummingbird (*Archilochus colubris*). The Auk 127, 626–635.

Berns, C.M., and Adams, D.C. (2013). Becoming different but staying alike: Patterns of sexual size and shape dimorphism in bills of hummingbirds. Evolutionary Biology 40, 246–260.

Birn-Jeffery, A.V., Miller, C.E., Naish, D., Rayfield, E.J., and Hone, D.W. (2012). Pedal claw curvature in birds, lizards and Mesozoic dinosaurs–complicated categories and compensating for mass-specific and phylogenetic control. PloS One 7, e50555.

Blüthgen, N., Menzel, F., and Blüthgen, N. (2006). Measuring specialization in species interaction networks. BMC Ecology 6, 9–21.

Boehm, M.M.A. (2018). Biting the hand that feeds you: Wedge-billed hummingbird is a nectar robber of a sicklebill-adapted andean bellflower. Acta Amazonica 48, 146–150.

Boehm, M.M.A. (2020). Curvr: Calculate Point-Wise Curvature From Landmarked Specimens.

Bonhomme, V., Picq, S., Gaucherel, C., and Claude, J. (2014). Momocs: Outline analysis using R. Journal of Statistical Software 56, 1–24.

Bookstein, F.L. (1991). Morphometric tools for landmark data: geometry and biology (Cambridge University Press).

Borcard, D., Gillet, F., and Legendre, P. (2018). Numerical Ecology with R (New York, USA: Springer).

Borchers, H.W. (2019). Pracma: Practical numerical math functions. R package version 2.2.5.

Borgella Jr, R., Snow, A.A., and Gavin, T.A. (2001). Species richness and pollen loads of hummingbirds using forest fragments in southern Costa Rica. Biotropica 33, 90–109.

Bright, J.A., Marugán-Lobón, J., Cobb, S.N., and Rayfield, E.J. (2016). The shapes of bird beaks are highly controlled by nondietary factors. Proceedings of the National Academy of Sciences 113, 5352–5357.

Campos, E.O., Bradshaw, H.D., and Daniel, T.L. (2015). Shape matters: Corolla curvature improves nectar discovery in the hawkmoth *Manduca sexta*. Functional Ecology 29, 462–468.

Cardinal, S., and Danforth, B.N. (2013). Bees diversified in the age of eudicots. Proceedings of the Royal Society B: Biological Sciences 280, 20122686.

Carothers, J.H. (1982). Effects of trophic morphology and behavior on foraging rates of three Hawaiian honeycreepers. Oecologia 55, 157–159.

Casey, J. (1996). Exploring Curvature (Braunschweig, Germany: Friedr. Vieweg & Sohn Verlagsgesellschaft mbH).

Castle, E.S. (1962). Phototropic curvature in *Phycomyces*. The Journal of General Physiology 45, 743–756.

Claude, J. (2008). Morphometrics with R (New York, USA: Springer Science).

Cleveland, W.S., Grosse, E., Shyu, W.M., Chambers, J.M., and Hastie, T.J. (1992). Statistical models in s. Local Regression Models Chapter–8.

Collins, B.G. (2008). Nectar intake and foraging efficiency: Responses of honeyeaters and hummingbirds to variations in floral environments. The Auk 125, 574–587.

Collyer, M.L., and Adams, D.C. (2018). RRPP: An R package for fitting linear models to high-dimensional data using residual randomization. Methods in Ecology and Evolution 9, 1772–1779.

Collyer, M., Sekora, D., and Adams, D.C. (2015). A method for analysis of phenotypic change for phenotypes described by high-dimensional data. Heredity 115, 357–365.

Coolidge, J.L. (1952). The unsatisfactory story of curvature. The American Mathematical Monthly 59, 375–379.

Cosgrove, D.J. (1990). Rapid, bilateral changes in growth rate and curvature during gravitropism of cucumber hypocotyls: Implications for mechanism of growth control. Plant, Cell & Environment 13, 227–234.

Cotton, P.A. (1998). Temporal partitioning of a floral resource by territorial hummingbirds. Ibis 140, 647–653.

Dakin, R., Segre, P.S., Straw, A.D., and Altshuler, D.L. (2018). Morphology, muscle capacity, skill, and maneuvering ability in hummingbirds. Science 359, 653–657.

Dehling, D.M., Töpfer, T., Schaefer, H.M., Jordano, P., Böhning-Gaese, K., and Schleuning, M. (2014). Functional relationships beyond species richness patterns: Trait matching in plant–bird mutualisms across scales. Global Ecology and Biogeography 23, 1085–1093.

Dellinger, A.S., Chartier, M., Fernández-Fernández, D., Penneys, D.S., Alvear, M., Almeda, F., Michelangeli, F.A., Staedler, Y., Armbruster, W.S., and Schönenberger, J. (2019). Beyond buzz–pollination-departures from an adaptive plateau lead to new pollination syndromes. New Phytologist 221, 1136–1149.

Emlen, D.J., Marangelo, J., Ball, B., and Cunningham, C.W. (2005). Diversity in the weapons of sexual selection: Horn evolution in the beetle genus *Onthophagus*. Evolution 59, 1060–1084.

Feinsinger, P., and Colwell, R.K. (1978). Community organization among neotropical nectar-feeding birds. American Zoologist 18, 779–795.

Frassen, J.M., Firn, R.D., and Digby, J. (1982). The role of the apex in the phototropic curvature of *Avena* coleoptiles: Positive curvature under conditions of continuous illumination. Planta 155, 281–286.

Gill, F.B., and Wolf, L.L. (1978). Comparative foraging efficiencies of some montane sunbirds in Kenya. The Condor 80, 391–400.

Goriely, A., and Neukirch, S. (2006). Mechanics of climbing and attachment in twining plants. Physical Review Letters 97, 184302.

Gómez, J., Perfectti, F., Bosch, J., and Camacho, J. (2009). A geographic selection mosaic in a generalized plant–pollinator–herbivore system. Ecological Monographs 79, 245–263.

Graham, C.H., Parra, J.L., Rahbek, C., and McGuire, J.A. (2009). Phylogenetic structure in tropical hummingbird communities. Proceedings of the National Academy of Sciences 106, 19673–19678.

Grant, V. (1949). Pollination systems as isolating mechanisms in angiosperms. Evolution 3, 82–97.

Grant, V., and Grant, K.A. (1983). Hawkmoth pollination of *Mirabilis longiflora* (Nyctaginaceae). Proceedings of the National Academy of Sciences 80, 1298–1299.

Hadley, A.S., Frey, S.J., Robinson, W.D., and Betts, M.G. (2018). Forest fragmentation and loss reduce richness, availability, and specialization in tropical hummingbird communities. Biotropica 50, 74–83.

Hainsworth, F.R. (1973). On the tongue of a hummingbird: Its role in the rate and energetics of feeding. Comparative Biochemistry and Physiology Part A: Physiology 46, 65–78.

Hughes, M., Möller, M., Edwards, T.J., Bellstedt, D.U., and De Villiers, M. (2007). The impact of pollination syndrome and habitat on gene flow: A comparative study of two *Streptocarpus* (Gesneriaceae) species. American Journal of Botany 94, 1688–1695.

Johnson, S.D., Kiepiel, I., and Robertson, A.W. (2020). Functional consequences of flower curvature, orientation and perch position for nectar feeding by sunbirds. Biological Journal of the Linnean Society 131, 822–834.

Joly, S., Lambert, F., Alexandre, H., Clavel, J., Léveillé-Bourret, É., and Clark, J.L. (2018). Greater pollination generalization is not associated with reduced constraints on corolla shape in Antillean plants. Evolution 72, 244–260.

Kay, K.M., and Sargent, R.D. (2009). The role of animal pollination in plant speciation: Integrating ecology, geography, and genetics. Annual Review of Ecology, Evolution, and Systematics 40, 637–656.

Klingenberg, C.P. (2016). Size, shape, and form: Concepts of allometry in geometric morphometrics. Development Genes and Evolution 226, 113–137.

Kuhl, F.P., and Giardina, C.R. (1982). Elliptic Fourier features of a closed contour. Computer Graphics and Image Processing 18, 236–258.

Lagomarsino, L.P., and Muchhala, N. (2019). A gradient of pollination specialization in three species of Bolivian *Centropogon*. American Journal of Botany 106, 633–642.

Lagomarsino, L.P., Forrestel, E.J., Muchhala, N., and Davis, C.C. (2017). Repeated evolution of vertebrate pollination syndromes in a recently diverged Andean plant clade. Evolution 71, 1970–1985.

Lenth, R., Singmann, H., Love, J., and others (2018). Emmeans: Estimated marginal means, aka least-squares means. R Package Version 1.

Luo, Y.-L., and Li, Q.-J. (2010). Effects of light and low temperature on the reciprocal style curvature of flexistylous Alpinia species (Zingiberaceae). Acta Physiologiae Plantarum 32, 1229–1234.

MacLeod, N. (1999). Generalizing and extending the eigenshape method of shape space visualization and analysis. Paleobiology 25, 107–138.

MacLeod, N. (2002). Geometric morphometrics and geological shape-classification systems. Earth-Science Reviews 59, 27–47.

MacLeod, N. (2011). The centre cannot hold I: ZR fourier analysis. Palaeontological Association Newsletter 78, 35–45.

MacLeod, N. (2012). The centre cannot hold II: Elliptic fourier analysis. Palaeontological Association Newsletter 79, 29–42.

MacLeod, N., and Rose, K.D. (1993). Inferring locomotor behavior in paleogene mammals via eigenshape analysis. American Journal of Science 293, 300–355.

Maglianesi, M.A., Blüthgen, N., Böhning-Gaese, K., and Schleuning, M. (2014). Morphological traits determine specialization and resource use in plant–hummingbird networks in the Neotropics. Ecology 95, 3325–3334.

Maglianesi, M.A., Böhning-Gaese, K., and Schleuning, M. (2015b). Different foraging preferences of hummingbirds on artificial and natural flowers reveal mechanisms structuring plant–pollinator interactions. Journal of Animal Ecology 84, 655–664.

Maglianesi, M.A., Blüthgen, N., Böhning-Gaese, K., and Schleuning, M. (2015a). Functional structure and specialization in three tropical plant–hummingbird interaction networks across an elevational gradient in Costa Rica. Ecography 38, 1119–1128.

Manning, J.C., and Goldblatt, P. (1997). The *Moegistorhynchus longirostris* (Diptera: Nemestrinidae) pollination guild: Long-tubed flowers and a specialized long-proboscid fly pollination system in southern Africa. Plant Systematics and Evolution 206, 51–69.

Martén-Rodríguez, S., Almarales-Castro, A., and Fenster, C.B. (2009). Evaluation of pollination syndromes in Antillean Gesneriaceae: Evidence for bat, hummingbird and generalized flowers. Journal of Ecology 97, 348–359.

Maruyama, P.K., Sonne, J., Vizentin-Bugoni, J., Martín González, A.M., Zanata, T.B., Abrahamczyk, S., Alarcón, R., Araujo, A.C., Araújo, F.P., Baquero, A.C., et al. (2018). Functional diversity mediates macroecological variation in plant–hummingbird interaction networks. Global Ecology and Biogeography 27, 1186–1199.

Mclntyre, G., and Browne, K. (1996). Effect of darkening the cotyledons on the growth and curvature of the sunflower hypocotyl: Evidence of hydraulic signalling. Journal of Experimental Botany 47, 1561–1566.

Miller, E.T., Wagner, S.K., Harmon, L.J., and Ricklefs, R.E. (2017). Radiating despite a lack of character: Ecological divergence among closely related, morphologically similar honeyeaters (Aves: Meliphagidae) co-occurring in arid Australian environments. The American Naturalist 189, E14–E30.

Milnor, J. (1954). On total curvatures of closed space curves. Mathematica Scandinavica 1, 289–296.

Minnaar, C., Jager, M. de, and Anderson, B. (2019). Intraspecific divergence in floral-tube length promotes asymmetric pollen movement and reproductive isolation. New Phytologist 1160–1170.

Mountainspring, S. (1987). Ecology, behavior, and conservation of the Maui Parrotbill. The Condor 89, 24–39.

Muller, A. (1995). Morphological specializations in Central European bees for the uptake of pollen from flowers with anthers hidden in narrow corolla tubes (Hymenoptera: Apoidea). Entomologia Generalis 20, 43–57.

Navalón, G., Marugán-Lobón, J., Bright, J.A., Cooney, C.R., and Rayfield, E.J. (2020). The consequences of craniofacial integration for the adaptive radiations of Darwin’s finches and Hawaiian honeycreepers. Nature Ecology & Evolution 4, 270–278.

Niet, T. van der, and Johnson, S.D. (2012). Phylogenetic evidence for pollinator-driven diversification of angiosperms. Trends in Ecology & Evolution 27, 353–361.

Ollerton, J. (2017). Pollinator diversity: Distribution, ecological function, and conservation. Annual Review of Ecology, Evolution, and Systematics 48, 353–376.

Olsen, A.M. (2017). Feeding ecology is the primary driver of beak shape diversification in waterfowl. Functional Ecology 31, 1985–1995.

Partida-Lara, R., Enriquez, P.L., Vazquez Perez, J.R., and Pineda Diez de Bonilla, E. (2018). Spatio-temporal structure of the taxonomic and functional diversity of hummingbirds at the biosphere reserve El Triunfo, Chiapas, Mexico. Ornitologia Neotropical 29, 37–50.

Paton, D., and Collins, B. (1989). Bills and tongues of nectar-feeding birds: A review of morphology, function and performance, with intercontinental comparisons. Australian Journal of Ecology 14, 473–506.

Peng, F., Campos, E.O., Sullivan, J.G., Berry, N., Song, B.B., Daniel, T.L., and Bradshaw Jr, H. (2019). Morphospace exploration reveals divergent fitness optima between plants and pollinators. PloS One 14, e0213029.

Pour, A.S., Chegini, G., Zarafshan, P., and Massah, J. (2018). Curvature-based pattern recognition for cultivar classification of *Anthurium* flowers. Postharvest Biology and Technology 139, 67–74.

R Core Team (2017). R: A language and environment for statistical computing (Vienna, Austria: R Foundation for Statistical Computing).

Raudseps, J.G. (1965). Some aspects of the tangent-angle vs. Arc length representation of contours (Ohio State University Research Foundation, Communication; Control Systems Lab).

Rico-Guevara, A., and Araya-Salas, M. (2014). Bills as daggers? A test for sexually dimorphic weapons in a lekking hummingbird. Behavioral Ecology 26, 21–29.

Ritter, R. (1992). Lateral bending during lizard locomotion. Journal of Experimental Biology 173, 1–10.

Robertson, C. (1889). Flowers and Insects. II. Botanical Gazette 14, 172–178.

Rocha, D., Monte Bello, C., Sobol, S., Samach, A., and Dornelas, M. (2015). Auxin and physical constraint exerted by the perianth promote androgynophore bending in *Passiflora mucronata* L. (Passifloraceae). Plant Biology 17, 639–646.

Rohlf, F.J. (1990). Fitting Curves to Outlines. In Proceedings of the Michigan Morphometrics Workshop, F.J. Rolhf, and F.L. Bookstein, eds. (Ann Arbor, MI: University of Michigan Museum of Zoology), pp. 177–188.

Rohlf, F.J. (2015). The tps series of software. Hystrix 26.

Rohlf, F., and Slice, D. (1990). Extensions of the Procrustes method for optimal superposition of landmarks. Systematic Zoology 39, 40–59.

Rutter, J.W. (2000). Geometry of Curves (Boca Raton, FL: CRC Press, Taylor; Francis Group).

Schmitt, D., Rose, M.D., Turnquist, J.E., and Lemelin, P. (2005). Role of the prehensile tail during ateline locomotion: Experimental and osteological evidence. American Journal of Physical Anthropology 126, 435–446.

Scott-Elliot, G. (1890). Ornithophilous flowers in South Africa. Annals of Botany 4, 265–280.

Silk, W.K. (1989). On the curving and twining of stems. Environmental and Experimental Botany 29, 95–109.

Smith, T.B., Freed, L.A., Lepson, J.K., and Carothers, J.H. (1995). Evolutionary consequences of extinctions in populations of a Hawaiian honeycreeper. Conservation Biology 9, 107–113.

Snow, B.K., and Snow, D. (1972). Feeding niches of hummingbirds in a Trinidad valley. The Journal of Animal Ecology 471–485.

Sonne, J., Zanata, T.B., Martín González, A.M., Cumbicus Torres, N.L., Fjeldså, J., Colwell, R.K., Tinoco, B.A., Rahbek, C., and Dalsgaard, B. (2019). The distributions of morphologically specialized hummingbirds coincide with floral trait matching across an Andean elevational gradient. Biotropica 51, 205–218.

Sonne, J., Vizentin-Bugoni, J., Maruyama, P.K., Araujo, A.C., Chávez-González, E., Coelho, A.G., Cotton, P.A., Marín-Gómez, O.H., Lara, C., Lasprilla, L.R., et al. (2020). Ecological mechanisms explaining interactions within plant–hummingbird networks: Morphological matching increases towards lower latitudes. Proceedings of the Royal Society B 287, 20192873.

Stearn, W.T. (2002). The genus Epimedium and other herbacious Berberidaceae including the genus Podophyllum (Portland, OR: Timber Press, Inc.).

Stiles, F.G. (1975). Ecology, flowering phenology, and hummingbird pollination of some Costa Rican *Heliconia* species. Ecology 56, 285–301.

Stiles, F.G. (1995). Behavioral, ecological and morphological correlates of foraging for arthropods by the hummingbirds of a tropical wet forest. Condor 853–878.

Stiles, F.G. (2004). Phylogenetic constraints upon morphological and ecological adaptation in hummingbirds (Trochilidae): Why are there no hermits in the paramo. Ornitologia Neotropical 15, 191–198.

Stiles, F.G. (2008). Ecomorphology and phylogeny of hummingbirds: Divergence and convergence in adaptations to high elevations. Ornitologia Neotropical 19, 511–519.

Temeles, E.J. (1996). A new dimension to hummingbird-flower relationships. Oecologia 105, 517–523.

Temeles, E.J., and Kress, W.J. (2003). Adaptation in a plant-hummingbird association. Science 300, 630–633.

Temeles, E.J., Pan, I.L., Brennan, J.L., and Horwitt, J.N. (2000). Evidence for ecological causation of sexual dimorphism in a hummingbird. Science 289, 441–443.

Temeles, E.J., Goldman, R.S., and Kudla, A.U. (2005). Foraging and territory economics of sexually dimorphic Purple-Throated Caribs (*Eulampis jugularis*) on three *Heliconia* morphs. The Auk 122, 187–204.

Temeles, E.J., Koulouris, C.R., Sander, S.E., and Kress, W.J. (2009). Effect of flower shape and size on foraging performance and trade-offs in a tropical hummingbird. Ecology 90, 1147–1161.

Temeles, E.J., Miller, J.S., and Rifkin, J.L. (2010). Evolution of sexual dimorphism in bill size and shape of hermit hummingbirds (Phaethornithinae): A role for ecological causation. Philosophical Transactions of the Royal Society B: Biological Sciences 365, 1053–1063.

Terral, J.-F., Alonso, N., Capdevila, R.B.i., Chatti, N., Fabre, L., Fiorentino, G., Marinval, P., Jordá, G.P., Pradat, B., and Rovira, N. (2004). Historical biogeography of olive domestication (*Olea europaea* L.) As revealed by geometrical morphometry applied to biological and archaeological material. Journal of Biogeography 31, 63–77.

Travers, S.E., Temeles, E.J., and Pan, I. (2003). The relationship between nectar spur curvature in jewelweed (*Impatiens capensis*) and pollen removal by hummingbird pollinators. Canadian Journal of Botany 81, 164–170.

Tripp, E.A., and McDade, L.A. (2013). Time-calibrated phylogenies of hummingbirds and hummingbird-pollinated plants reject a hypothesis of diffuse co-evolution. Journal of Systematic and Evolutionary Botany 31, 89–103.

Vamosi, J.C., Magallón, S., Mayrose, I., Otto, S.P., and Sauquet, H. (2018). Macroevolutionary patterns of flowering plant speciation and extinction. Annual Review of Plant Biology 69, 685–706.

Van Otterloo, P.J. (1991). A contour-oriented approach to shape analysis (Prentice Hall New York, NY).

Wang, Q., Li, Y., Pu, X., Zhu, L., Tang, Z., and Liu, Q. (2013). Pollinators and nectar robbers cause directional selection for large spur circle in *Impatiens oxyanthera* (Balsaminaceae). Plant Systematics and Evolution 299, 1263–1274.

Webster, M., and Sheets, H.D. (2010). A practical introduction to landmark-based geometric morphometrics. The Paleontological Society Papers 16, 163–188.

Weinstein, B.G., and Graham, C.H. (2017). Persistent bill and corolla matching despite shifting temporal resources in tropical hummingbird-plant interactions. Ecology Letters 20, 326–335.

Wood, S.N. (2017). Generalized additive models: An introduction with r (Boca Raton, FL: CRC Press).

Young, H.J. (2008). Selection on spur shape in *Impatiens capensis*. Oecologia 156, 535–543.

Zahn, C.T., and Roskies, R.Z. (1972). Fourier descriptors for plane closed curves. IEEE Transactions on Computers 100, 269–281.

Zwillinger, D. (2018). In CRC Standard Mathematical Tables and Formulae, (Boca Raton, FL: Chapman; Hall, CRC Press), pp. 424–425.

