## Supplementary Materials for "Plant-pollinator specialization: Origin and measurement of curvature"

### Supplementary material for Plant-pollinator specialization: Origin and measurement of curvature

#### Measuring *Epimedium* flowers and defining developmental stages

We tested the utility of the proposed curvature metric by studying floral development in *Epimedium* L. (Berberidaceae), a group of temperate, perennial herbs that inhabit montane ecosystems from North Africa to East Asia. *Epimedium* is perhaps best recognized by having an elongated, curved nectar spur - a derived trait for pollination by bees (Stearn, 2002).

Although widespread throughout *Epimedium*, the ecological function of nectar spur curvature is largely unstudied. Early work comparing sympatric short- and long-spurred species found evidence for pollinator partitioning in Japanese *Epimedium* (Suzuki, 1984): long-horned bees with short proboscides (*Eucerea nipponensis* Pérez) tend to visit *Epimedium trifoliolatobinatum* Koidz (spur length 10-15 mm), while bumblebees with longer proboscides ( *Bombus diversus* Smith) visit *Epimedium grandiflorum* C.Morren (spur length 10-20 mm. While these pollinators are nectar foraging, Suzuki (1984) also found that pollen-collecting bees (*Andrena* Fabricius and *Lassioglossum* Curtis) visit *Epimedium* indiscriminately, potentially contributing to hybridization. Although numerous *Epimedium* spp. exhibit strong floral curvature, the role of curvature, if any, on pollinator partitioning is unknown.

Flower size of *E. koreanum* and *E. violaceum* ($n$=83) was measured daily from April 9 to May 2, 2019 at the UBC Botanical Garden (TableS2). Size was defined as the width between the apex of the two outer sepals on the major axis of the flower (aestivation is imbricate). Width was measured to the nearest 0.1 mm using an SPI Polymid Dial Caliper. By correlating changes in flower size to developmental progress (Figure S3), we were able to define 4 discrete size-stages present in both taxa (Table S3, Figure S4).

In *E. violaceum* the distance sepal distance was measured until the width of the inner sepals exceeded the width of the outer sepals (‘G’ Stage). From this point onwards, the inner sepal distance was measured. In *E. koreanum*, the inner sepals lack pigmentation and adhere closely to the petals, making them difficult to measure accurately. For this reason, the outer sepals were measured until they abscised (“T” stage).

Changes in flower size were punctuated by developmental milestones (defined in Table S3). We tested for the validity of these developmental stages by fitting a linear mixed effects model using the R package `lmerTest` v.3.1-2 (Kuznetsova et al., 2017):

$$sepal\ \ size = \beta_0 + \beta_1 * stage * taxon + U_i + \epsilon$$

Where $\beta_0$ is the intercept, $\beta_1$ is a coefficient, $U_i$ is the individual-specific random effect, and $\epsilon$ is the residual error. T-values were approximated using the Welch–Satterthwaite method in `lmerTest`. All data herein was organized and plotted using the R packages contained in the `tidyverse` (**???**)

We then tested for pairwise differences of estimated marginal means using the R package `emmeans` (Lenth et al., 2018). By correlating changes in flower size to developmental landmarks, we were able to define 4 discrete stages in *E. koreanum* and *E. violaceum* (Figure 5, Table S3). In the first stage (“C”) the petals are shorter in length than the sepals that envelop them - the following stage begins when the petals overtake the surrounding sepals in length. The ‘G’ stage includes continued growth of the bud until the petals begin to separate. At the ‘T’ stage nectar begins accumilating in the spurs. Anthesis takes place during the ‘A’ stage at which point the flower opening may increase in size and anthers dehisce.

Flowering stage data was staggered because each flower developed independently. The data was also fragmented because some samples abscissed prematurely due to herbivory or weather. The flower development data was therefore a ‘censored’ dataset i.e. some subjects left the study before reaching maturity. Although manually aligning the stage data is possible, for convenience we used a multiple sequence alignment protocol to automate the process. To do this, we ran the stage data through ClustalW implemented in the R package `msa` v.3.9 (Bodenhofer et al., 2015) with a neutral (identity) substitution matrix. Gap opening was prohibited. By aligning phenological data within species, a consensus (mean) stage sequence was calculated and used to estimate flower age where observations were censored.

We tested for differences in developmental time by fitting a linear mixed effects model using the R package `lmerTest`:

$$elpased\ \ days = \beta_0 + \beta_1 * stage * taxon + U_i + \epsilon$$

Where $\beta_0$ is the intercept, $\beta_1$ is a coefficient, $U_i$ is the individual-specific random effect, and $\epsilon$ is the residual error. We then tested for pairwise differences of estimated marginal means using the R package `emmeans`. Results are presented in Table S4.

#### Landmarking, morphometrics, and curve-fitting

Following the initial study of *Epimedium* development, a separate set of flowers were sampled for shape analysis (Table S2). Samples were first preserved in 70% ethanol and later transferred to a glass slide. Preserved flowers were imaged in the dorsiventral view using a stereo microscope at 6.3x (Zeiss Stemi 508 with Axiocam 301). Scale bars were automatically generated by the Zeiss Zen software interfacing with the camera. Specimens that did not fit within the field of view were imaged in halves and the images joined using the Stitching Plugin in the Fiji distribution of ImageJ2 (Preibisch et al., 2009; Rueden et al., 2017). To assign landmarks and semi-landmarks to the digitized specimens, we adhered to the following protocol:

1. Rotate the photographs so that the opening of the corolla tube is parallel to the y-axis.
2. Build .tps file (a file listing all specimens) using *tpsUtil* (Rohlf, 2015). This .tps file is used by *tpsdig* for landmark assignment.
3. Landmark specimens from .tps file using tpsDig (steps 1 and 2 can also be done in the R packages `MomX` (Bonhomme et al., 2014) and `geomorph` (Adams and Otárola-Castillo, 2013)). Landmarks used to measure the dorsal arc were (A) the farthest point on the apex of the spur before the inflection point where either the spur diminishes to a tip (*E. violaceum*) or widens into a saccate reservoir (*E. koreanum*), and (B) the dorsal point at which the spur widens to become an attachment point for the petal to the stem. 13 semi-landmarks (sensu Webster and Sheets, 2010) were placed between them (15 points total). Landmarks used to measure the ventral are (C) the analagous point to (A) on the ventral side of the spur, and (D) the crease created by the sudden widening of the narrow spur into a wide opening for pollinators. 13 semi-landmarks were placed between them (15 points total). An additional

landmark was place at the apex of the nectar spur, resulting in 31 total landmarks. The complete set of 31 landmarks was used for geometric morphometrics, whereas the dorsal and ventral 'sets' were used separately for curve-fitting and analysis of curvature.

4. Curve points are drawn in tpsDig using the "pencil tool" in tpsDig, from landmarks (A) to (B). Following the placement of points, a curve is automatically drawn that connects them. We used the *Resample Curve* function and selected to space the points evenly *by length*. We then manually adjusted the re-sampled points back onto the specimen and used the *Resample Curve* function until all landmarks were aligned and evenly spaced. This does not usually need to be repeated more than twice.
5. We set the scale by navigating *Options* -> *Image Tools* and typing in desired length and units. We selected *Set scale* and selected both ends of the scale bar (included in our images). We then navigated to back to the *Image options* dialogue box to confirm 'OK'.
6. Semi-landmarks need to be treated like landmarks for curve-fitting. To do this, we used the *Append tps curve to landmarks* function in tpsUtil.
7. Import appended .tps files into R using `from_tps()` function from `Momit` (Bonhomme et al., 2014) or `readmulti.tps()` from `geomorph`.

We used the R package `geomorph` (Adams and Otárola-Castillo, 2013) to study morphological development. First, landmark configurations (shapes) were aligned using a Generalized Procrustes Analysis (see: main text). Aligned landmark configurations (shapes) were then analysed for principal components desribing the major axes of shape variation. In `geomorph` this is done using `plotTangentSpace()`. We then fit a linear mixed model to correlate changes in shape with developmental stage:

$$shape = \beta_0 + \beta_1 * stage * taxon + U_i + \epsilon$$

Where $\beta_0$ is the intercept and $\beta_1$ is a coefficient, $U_i$ is the individual-specific random effect, and $\epsilon$ is the residual error. Summary statistics are presented in Table S5. We then visualized the developmental trajectory through shape space for each taxon by implementing `geomorph::trajectory.analysis()` (Figure 4).

To investigate whether variation in shape can be attributed to curvature, we fit smoothing cubic splines to landmark coordinates for each specimen using `stats::smooth.spline()`. Shapes were pre-aligned so that the distance between landmarks (A) and (B) was parallel with the x-axis. The use of splines in modelling open curves is discussed in Rohlf (1990). Arc length was calculated from bounded polynomial functions using `pracma` v.2.2.5 (Borchers, 2019). Curvature, as defined in the previous section, was computed using custom functions available as an R package `curvr` hosted at github.com/mannfred/curvr. We tested for differences in the development of curvature by fitting a linear model:

$$curvature = \beta_0 + \beta_1 * stage * taxon + \epsilon$$

Where $\beta_0$ is the intercept, $\beta_1$ is a coefficient, and $\epsilon$ is the residual error. Because the covariance matrix of the of the random effect was singular (i.e. variance estimate was nearly zero), the variable $U_i$ was omitted (see previous equation). We then tested for pairwise differences of means using the `stats::TukeyHSD()` function. Results are presented in Table S6.

#### Supplementary Figures

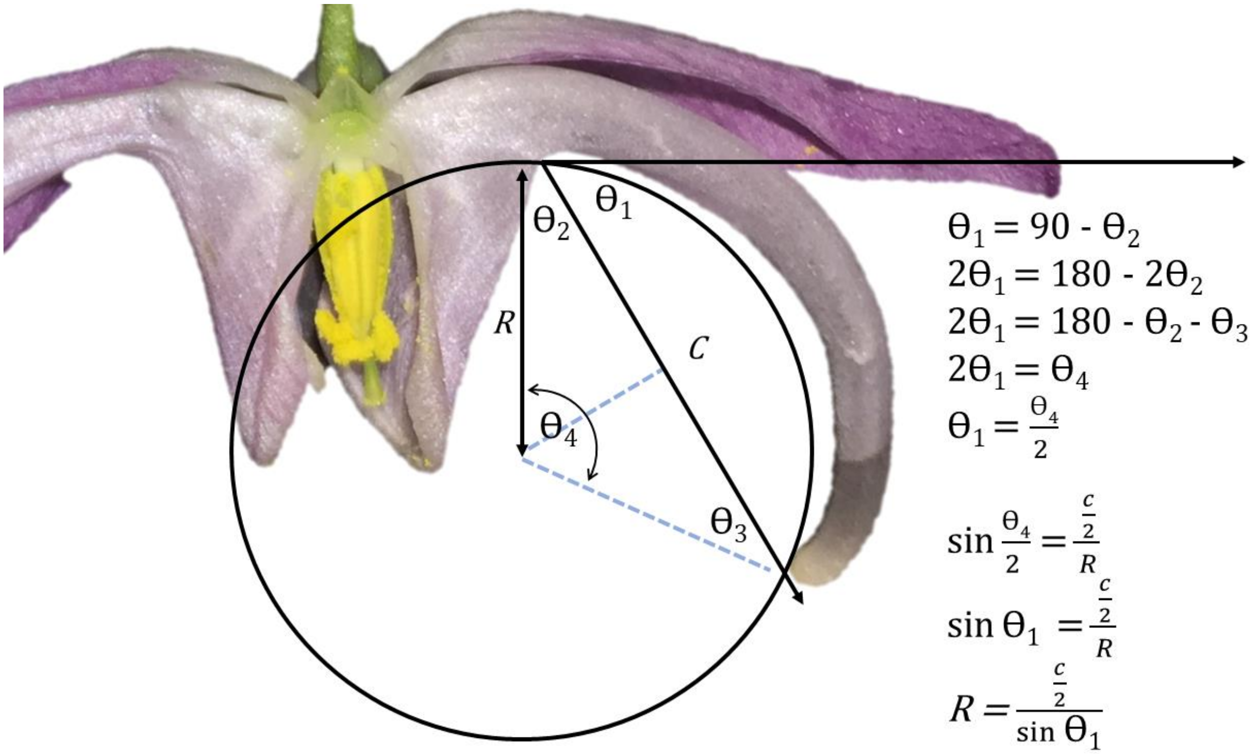


*Figure S1. Demonstration that the angle of deflection and inverse radius methods are interchangeable.*

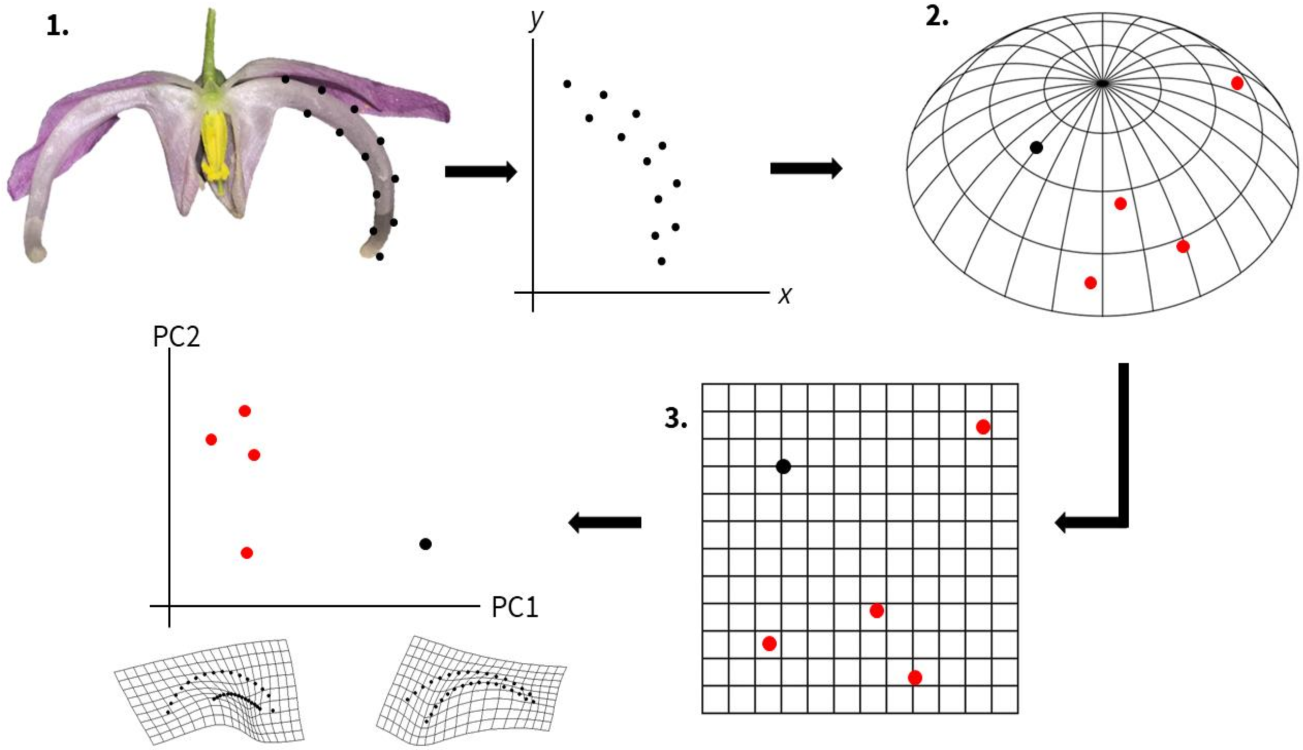


*Figure S2. Overview of a geometric morphometrics protocol. 1. Landmarks and semi-landmarks are assigned to a specimen. Each landmark is assigned an xy coordinate. 2. For each specimen a configuration of landmarks exists as a single point in a non-Euclidian shape space (abstracted here as a sphere segment). Red points represent landmark configurations from other specimens. 3. Shape data is projected onto a Euclidian plane – a tangent space approximation. This allows statistical analyses of shape variation (e.g. principal components analysis).*

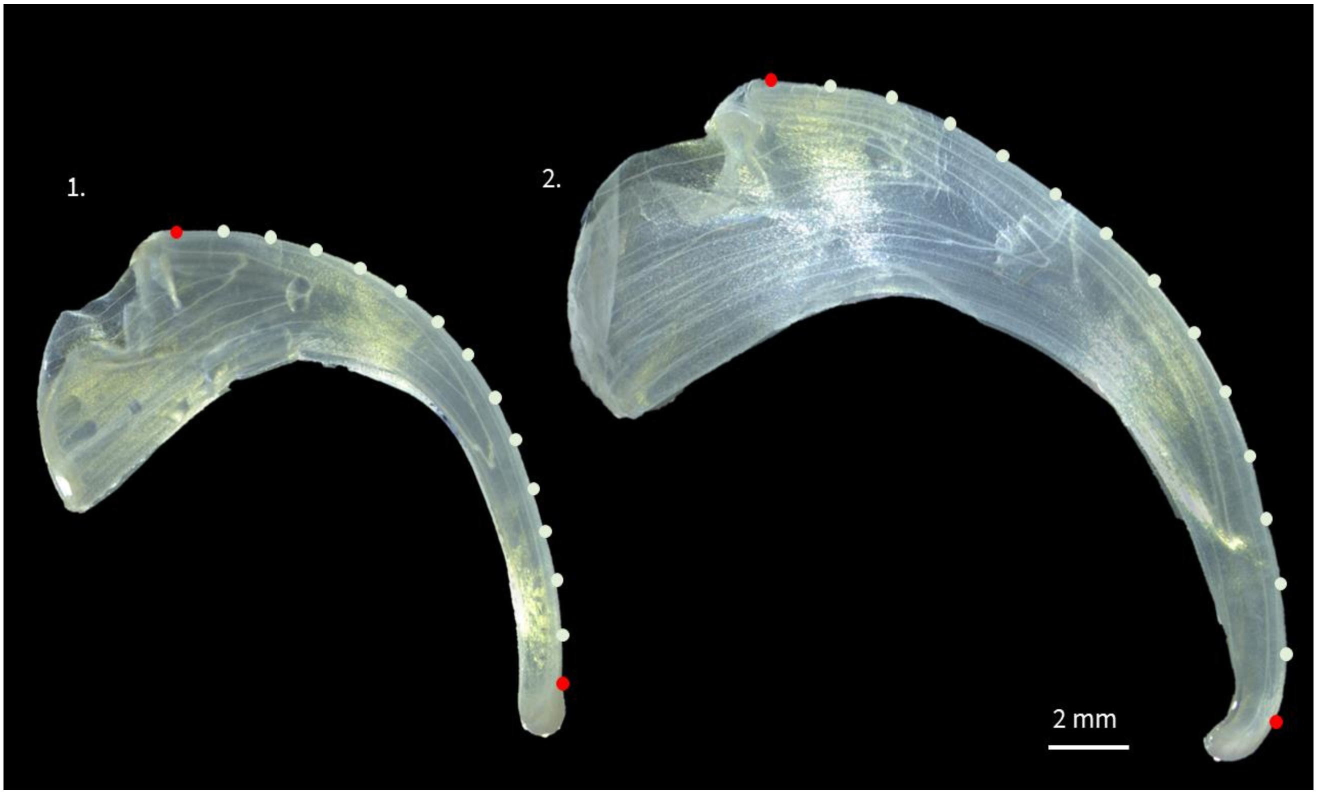


*Figure S3. Landmarks (red) and semi-landmarks (white) used to compare curvature between E. koreanum (left) and E. violaceum (right). Landmark #1 is designated at the apex of the nectar spur. Landmark #15 is placed at the base of the nectar spur. Petals sampled at anthesis.*

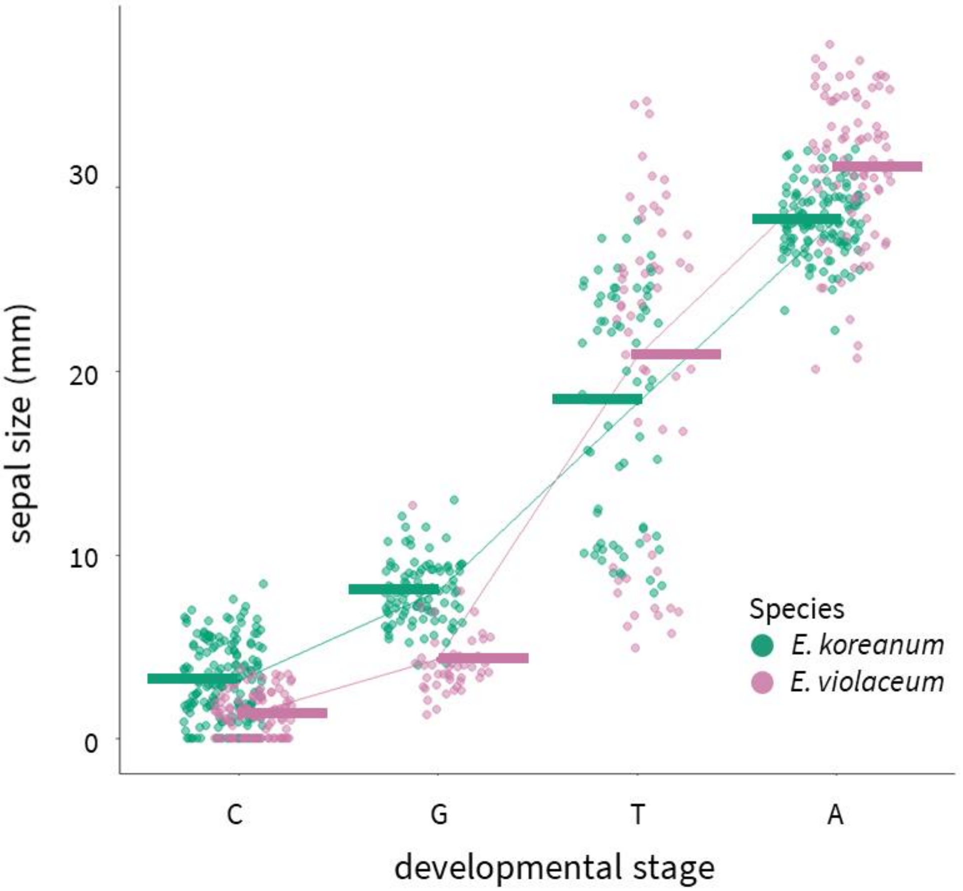


*Figure S4: Comparison of developmental stages and size in Epimedium. Tukey's HSD: p<0.01 for all within-species comparisons. Horizontal bars are least-squares means.*

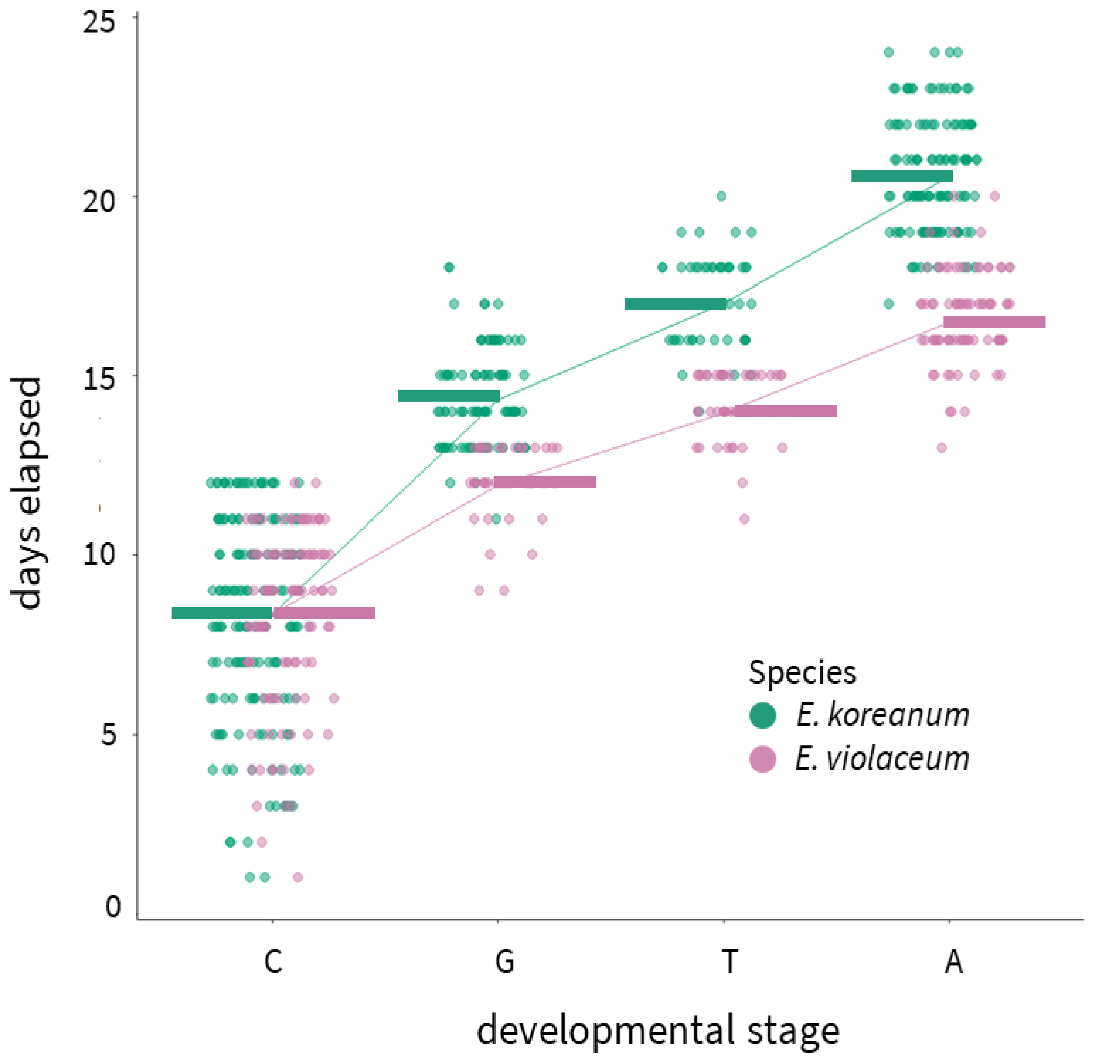


*Figure S5: Comparison of timing of developmental stages in Epimedium. Horizontal bars are least-squares means.*

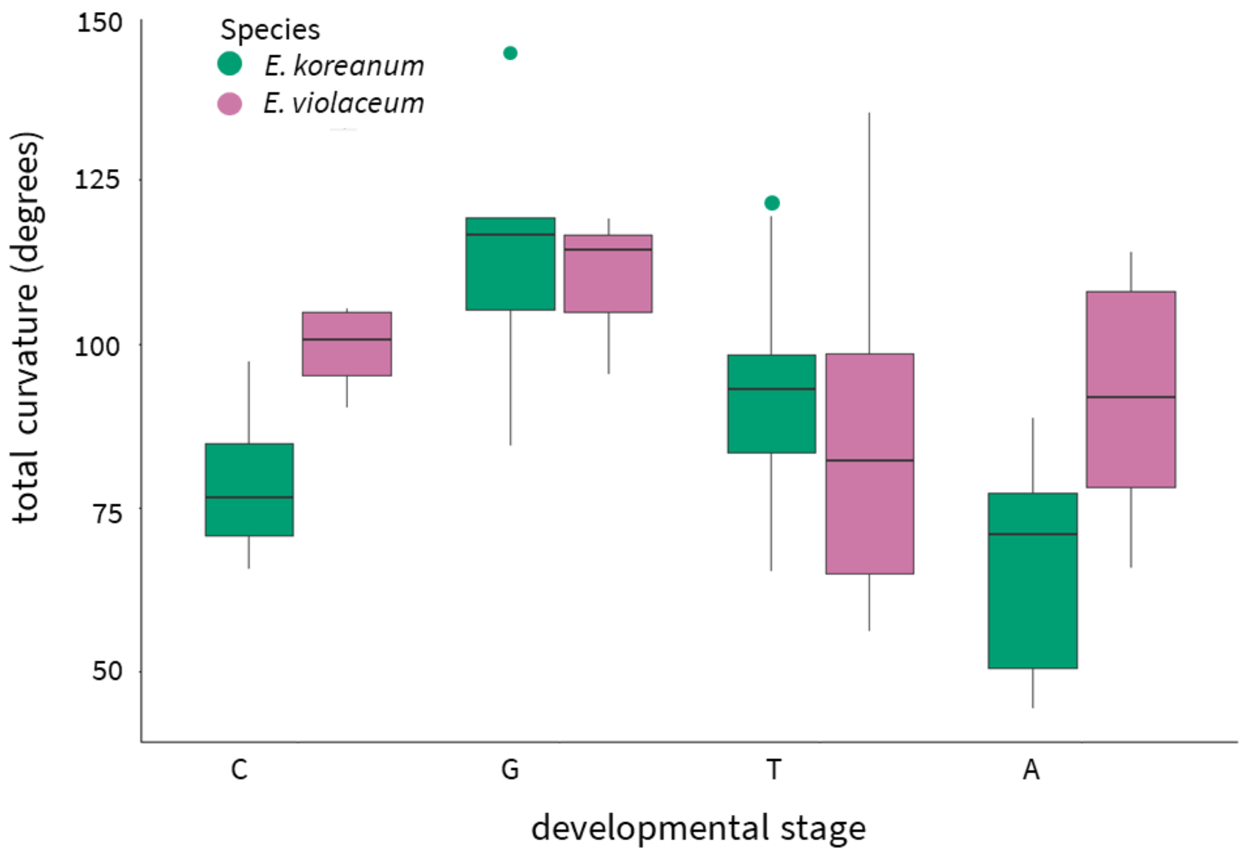


*Figure S6: Total curvature as a function of developmental stage in Epimedium.*

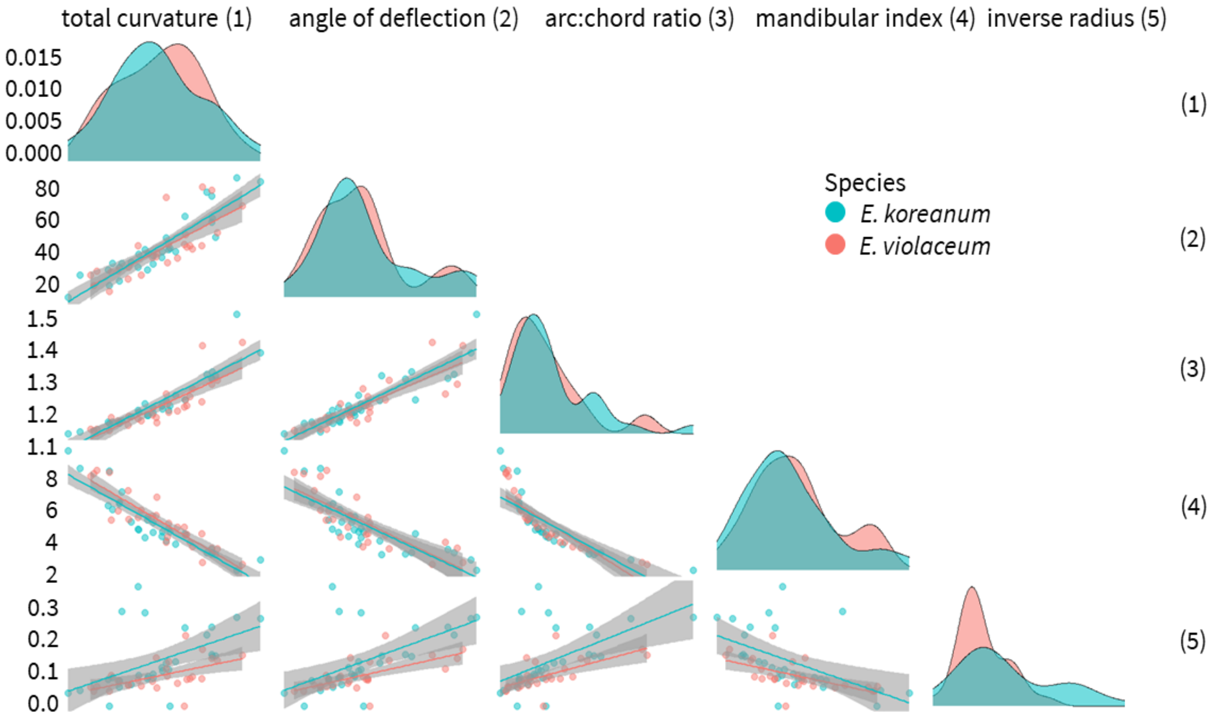


*Figure S7: Pairwise comparisons of curvature metrics. E. koreanum is red, E. violaceum is blue. Density plots show the distribution of curvature values for each species.*

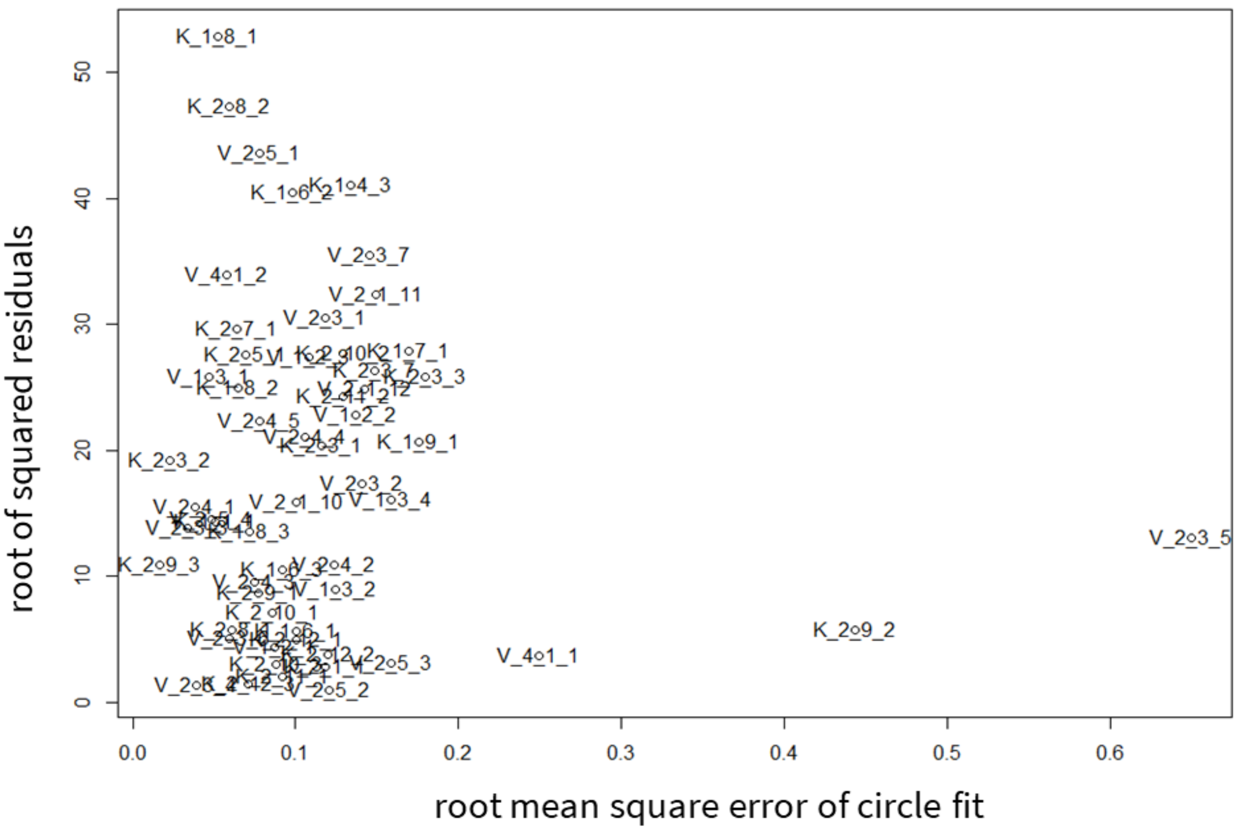


Figure S8: Scatterplot of x: the error distance for each specimen fitted by a circle - lower values are better approximated by a circle, and y: the residual distance between the inverse radius metric and total curvature - greater values for samples where the two metrics disagreed. Samples labeled with 'K' belong to E. koreanum. Samples labeled with 'V' belong to E. violaceum.

#### Supplementary Tables

166 *Table S1: Additional literature reviewed for metrics of floral or mouthpart curvature outside of*
167 *plant-pollinator systems.*

| Citation | System | Stated or Inferred Method |
|---|---|---|
| Baldwin et al. (1931) | Methods for univariate measurements and traditional morphometrics of birds. | inverse radius |
| Hamilton (1975) | Feeding ecology and behaviour of the American Avocet and the Black-necked Stilt (Recurvirostridae). | inverse radius |
| Buttrose et al. (1977) | Style curvature as a mechanism for self-pollination in *Hibiscus trionum* (Malvaceae). | qualitative |
| Ortiz et al. (2000) | Development of corolla curvature in herkogamous *Putoria calabrica* (Rubiaceae). | qualitative |
| Lindqvist et al. (2003) | Cladistics, taxonomy, and floral diversity of mints (Lamiaceae) endemic to Hawaii. | qualitative |
| Ruan et al. (2008) | Style curvature in autogamous *Kosteletzkya virginica* (Malvaceae). | qualitative |
| Kawabata et al. (2009) | Cultivar classification of *Eustoma* (Gentianaceae). | fourier analysis |
| Dalayap et al. (2011) | Varietal classification of Mokara orchids (Orchidaceae). | geometric morphometrics |

| | | |
|---|---|---|
| Nii and Kawabata (2011) | Cultivar classification of *Eustoma* (Gentianaceae). | fourier analysis |
| Berger et al. (2017) | Quantifying morphological modifications to floral form in gene knockdowns of *Fedia graciliflora* (Caprifoliaceae) | geometric morphometrics |
| Pour et al. (2018) | Cultivar classification of *Anthurium* (Araceae). | point-wise curvature |
| Song et al. (2018) | Cultivar classification of *Chrysanthemum* (Asteraceae). | angle of deflection |

*Table S2: Sample sizes for (A) flowers used to define developmental stages and (B) flowers used to quantify shape variation*

| Species | *n* (Individuals) | *n* (total panicles) | *n* (total flowers) |
|---|---|---|---|
| *E. koreanum* (A) | 2 | 11 | 50 |
| *E. violaceum* (A) | 2 | 3 | 33 |
| *E. koreanum* (B) | 2 | 15 | 30 |
| *E. violaceum* (B) | 2 | 7 | 27 |

*Table S3: Stages of Epimedium flower development. EMM is estimated maringal mean, SE is standard error, DF is degrees of freedom.*

| Stage | Defintion | Taxon | EMM (sepal size in mm) | SE | DF | Lower CL | Upper CL |
|---|---|---|---|---|---|---|---|
| C | Petals do not exceed the length of the inner and outer sepals. | *E. koreanum* | 3.21 | 0.479 | 3.18 | 0.128 | 6.30 |
| | | *E. violaceum* | 1.24 | 0.520 | 4.40 | 0.000 | 3.78 |
| G | Petals exceed the length of the inner and outer sepals. | *E. koreanum* | 8.09 | 0.511 | 4.14 | 5.493 | 10.70 |
| | | *E. violaceum* | 4.31 | 0.657 | 11.16 | 2.113 | 6.51 |
| T | Opening and separation of the petals. At least one petal is free from touching adjacent petals. Outer sepals begin to abscise. Nectar is visibly collecting in spurs. | *E. koreanum* | 18.19 | 0.542 | 5.22 | 15.812 | 20.57 |

| | | | | | | | |
|---|---|---|---|---|---|---|---|
| | | *E. violaceum* | 20.88 | 0.625 | 8.86 | 18.659 | 23.10 |
| A | Initiated by partial anther dehiscence, followed by complete dehiscence, and finally flower abscisson | *E. koreanum* | 27.84 | 0.476 | 3.10 | 24.694 | 30.99 |
| | | *E. violaceum* | 31.75 | 0.511 | 3.98 | 29.067 | 34.43 |

*Table S4: Timing of flower development in Epimedium. EMM is estimated maringal mean, SE is standard error, DF is degrees of freedom.*

| Developmental stage | Taxon | EMM (Elapsed days) | SE | DF | Lower CL | Upper CL |
|---|---|---|---|---|---|---|
| C | *E. koreanum* | 8.347212 | 0.3299095 | 2.507147 | 5.515885 | 11.17854 |
| | *E. violaceum* | 8.357769 | 0.3467908 | 3.058148 | 6.035077 | 10.68046 |
| G | *E. koreanum* | 14.313171 | 0.3512249 | 3.229466 | 12.092260 | 16.53409 |
| | *E. violaceum* | 11.941145 | 0.4080819 | 5.868882 | 10.252550 | 13.62974 |
| T | *E. koreanum* | 17.027220 | 0.3763897 | 4.227753 | 15.138820 | 18.91562 |
| | *E. violaceum* | 13.964860 | 0.3962915 | 5.145850 | 12.209970 | 15.71975 |
| A | *E. koreanum* | 20.559784 | 0.3422595 | 2.871271 | 18.096690 | 23.02288 |
| | *E. violaceum* | 16.498930 | 0.3657876 | 3.724022 | 14.472770 | 18.52509 |

*Table S5: Pairwise comparisons of shape through flower development in Epimedium. d is standardized (Cohen) effect size, UCL is upper confidence limit, Z is effect size adjusted by standard error. Contrasts are denoted by "Stage, Taxon" where K is E. koreanum and V is E. violaceum. Stages (young to old: C,G,T,A) defined in Table S3. Interspecific contrasts at the same stage are indicate by bold face. The strongest interspecific difference is at the earliest stage (C).*

| Constrast | d | UCL (95%) | Z | Pr > d |
|---|---|---|---|---|
| C,K - G,K | 0.219149 | 0.253073 | 0.723800 | 0.233 |
| C,K - T,K | 0.364526 | 0.307599 | 3.428385 | 0.001 |
| A,K - C,K | 0.383335 | 0.363423 | 2.191239 | 0.012 |
| **C,K - C,V** | **0.165909** | **0.116206** | **3.947645** | **0.002** |
| C,K - G,V | 0.241420 | 0.241740 | 1.737283 | 0.053 |
| C,K - T,V | 0.284976 | 0.295396 | 1.296141 | 0.098 |
| A,V - C,K | 0.332926 | 0.315041 | 2.173542 | 0.019 |
| G,K - T,K | 0.243366 | 0.259229 | 1.063465 | 0.140 |
| A,K - G,K | 0.298889 | 0.332143 | 0.800440 | 0.213 |
| C,V - G,K | 0.172273 | 0.254521 | -0.880430 | 0.807 |
| **G,K - G,V** | **0.060242** | **0.130639** | **-0.772680** | **0.760** |
| G,K - T,V | 0.196549 | 0.256767 | -0.270460 | 0.604 |
| A,V - G,K | 0.233378 | 0.294272 | -0.092170 | 0.543 |

| | | | | |
|---|---|---|---|---|
| A,K - T,K | 0.080747 | 0.119570 | 0.288666 | 0.380 |
| C,V - T,K | 0.212939 | 0.318102 | -1.749760 | 0.963 |
| G,V - T,K | 0.200922 | 0.281237 | -0.479270 | 0.666 |
| **T,K - T,V** | **0.091004** | **0.076195** | **3.006631** | **0.010** |
| A,V - T,K | 0.059356 | 0.098435 | -0.424110 | 0.605 |
| A,K - C,V | 0.228996 | 0.366537 | -2.039080 | 0.980 |
| A,K - G,V | 0.255293 | 0.347963 | -0.362320 | 0.643 |
| A,K - T,V | 0.113821 | 0.130961 | 1.079410 | 0.154 |
| **A,K - A,V** | **0.080805** | **0.115648** | **0.253307** | **0.371** |
| C,V - G,V | 0.157372 | 0.244312 | -0.569910 | 0.708 |
| C,V - T,V | 0.132221 | 0.304499 | -3.659050 | 1.000 |
| A,V - C,V | 0.176293 | 0.321780 | -2.547680 | 0.996 |
| G,V - T,V | 0.153234 | 0.273235 | -1.283770 | 0.896 |
| A,V - G,V | 0.187792 | 0.305365 | -1.052980 | 0.847 |
| A,V - T,V | 0.050740 | 0.094155 | -0.156720 | 0.474 |

*Table S6: Estimated means of total curvature through flower development in Epimedium. Contrasts are denoted by "Stage, Taxon" where K is E. koreanum and V is E. violaceum. Stages (young to old: C,G,T,A) defined in Table S3. Interspecific contrasts at the same stage are indicate by bold face.*

| contrast | difference in means | lower boundary | upper boundary | p adjusted |
|---|---|---|---|---|
| C,K - A,K | 12.677820 | -28.81440 | 54.170070 | 0.976967 |
| G,K - A,K | 50.613260 | 13.15941 | 88.067100 | 0.002076 |
| T,K - A,K | 25.904880 | -6.03587 | 57.845620 | 0.192882 |
| **A,V - A,K** | **25.049960** | **-10.21170** | **60.311600** | **0.342108** |
| C,V - A,K | 32.907970 | -8.58428 | 74.400210 | 0.215361 |
| G,V - A,K | 43.206230 | -1.96481 | 88.377270 | 0.070143 |
| T,V - A,K | 19.478250 | -13.44550 | 52.402020 | 0.575425 |
| G,K - C,K | 37.935440 | -1.99050 | 77.861370 | 0.073606 |
| T,K - C,K | 13.227050 | -21.57960 | 48.033680 | 0.927223 |
| A,V - C,K | 12.372140 | -25.50490 | 50.249210 | 0.966751 |
| **C,V - C,K** | **20.230140** | **-23.50650** | **63.966820** | **0.822150** |
| G,V - C,K | 30.528410 | -16.71260 | 77.769410 | 0.463271 |
| T,V - C,K | 6.800430 | -28.91040 | 42.511270 | 0.998687 |
| T,K - G,K | -24.708400 | -54.58620 | 5.169453 | 0.174358 |
| A,V - G,K | -25.563300 | -58.96770 | 7.841142 | 0.253853 |

| | | | | |
|---|---|---|---|---|
| C,V - G,K | -17.705300 | -57.63120 | 22.220640 | 0.851185 |
| **G,V - G,K** | **-7.407030** | **-51.14370** | **36.329640** | **0.999386** |
| T,V - G,K | -31.135000 | -62.06150 | -0.208510 | 0.047381 |
| A,V - T,K | -0.854910 | -27.93400 | 26.224180 | 1.000000 |
| C,V - T,K | 7.003091 | -27.80350 | 41.809710 | 0.998134 |
| G,V - T,K | 17.301350 | -21.81790 | 56.420620 | 0.852892 |
| **T,V - T,K** | **-6.426620** | **-30.38220** | **17.528940** | **0.989088** |
| C,V - A,V | 7.858001 | -30.01910 | 45.735070 | 0.997735 |
| G,V - A,V | 18.156260 | -23.71840 | 60.030940 | 0.865119 |
| T,V - A,V | -5.571710 | -33.80360 | 22.660190 | 0.998350 |
| G,V - C,V | 10.298260 | -36.94270 | 57.539260 | 0.996908 |
| T,V - C,V | -13.429700 | -49.14060 | 22.281130 | 0.930885 |
| T,V - G,V | -23.728000 | -63.65390 | 16.197960 | 0.569846 |

189

*Table S7: ANOVA for total curvature and PC2 of shape space. SE is standard error, df is degrees of freedom.*

| Model: PC2 ~ curvature*taxon | Estimate | SE | df | t value | Pr(>\|t\|) |
|---|---|---|---|---|---|
| intercept | -0.253890 | 0.034326 | 35.49263 | -7.396370 | 0.000000 |
| total curvature | 0.002933 | 0.000345 | 52.41149 | 8.492053 | 0.000000 |
| individual | -0.010360 | 0.051931 | 44.14297 | -0.199430 | 0.842842 |
| total curvature*taxon | -0.000140 | 0.000530 | 52.29149 | -0.267290 | 0.790297 |

*Table S8: Pairwise comparisons between historic metrics and total curvature for E. violaceum. Lower diagonal contains Pearson regression coefficients, upper diagonal contains p-values*

| | total curvature | angle of deflection | arc:chord ratio | mandibular index | 1/radius |
|---|---|---|---|---|---|
| total curvature | | 0.0000020 | 0.0000000 | 0.0000000 | 0.0030058 |
| angle of deflection | 0.7760905 | | 0.0000000 | 0.0000005 | 0.0002274 |
| arc:chord ratio | 0.8389153 | 0.8494683 | | 0.0000000 | 0.0003083 |
| mandibular index | -0.8951949 | -0.8000778 | -0.8636072 | | 0.0007902 |
| 1/radius | 0.5492380 | 0.6522140 | 0.6417751 | -0.6068571 | |

*Table S9: Pairwise comparisons between historic metrics and total curvature for E. koreanum. Lower diagonal contains Pearson regression coefficients, upper diagonal contains p-values*

| | total curvature | angle of deflection | arc:chord ratio | mandibular index | 1/radius |
|---|---|---|---|---|---|
| total curvature | | 0.0000000 | 0.0000000 | 0.0000000 | 0.0043597 |
| angle of deflection | 0.9126844 | | 0.0000000 | 0.0000000 | 0.0007646 |
| arc:chord ratio | 0.9046636 | 0.9075880 | | 0.0000001 | 0.0018090 |
| mandibular index | -0.9119176 | -0.8147521 | -0.8119942 | | 0.0031862 |
| 1/radius | 0.5057093 | 0.5808172 | 0.5458243 | -0.5205723 | |

### References


Adams, D.C., and Otárola-Castillo, E. (2013). Geomorph: An R package for the collection and analysis of geometric morphometric shape data. Methods in Ecology and Evolution *4*, 393–399.

Baldwin, S.P., Oberholser, H.C., and Worley, L.G. (1931). Measurements of birds (Cleveland Museum of Natural History).

Berger, B.A., Ricigliano, V.A., Savriama, Y., Lim, A., Thompson, V., and Howarth, D.G. (2017). Geometric morphometrics reveals shifts in flower shape symmetry and size following gene knockdown of *CYCLOIDEA* and *ANTHOCYANIDIN SYNTHASE*. BMC Plant Biology *17*, 205–214.

Bodenhofer, U., Bonatesta, E., Horejš-Kainrath, C., and Hochreiter, S. (2015). Msa: An r package for multiple sequence alignment. Bioinformatics *31*, 3997–3999.

Bonhomme, V., Picq, S., Gaucherel, C., and Claude, J. (2014). Momocs: Outline analysis using R. Journal of Statistical Software *56*, 1–24.

Borchers, H.W. (2019). Pracma: Practical numerical math functions. R package version 2.2.5.

Buttrose, M., Grant, W., and Lott, J. (1977). Reversible curvature of style branches of *Hibiscus trionum* L., a pollination mechanism. Australian Journal of Botany *25*, 567–570.

Dalayap, R.M., Torres, M.A.J., and Demayo, C.G. (2011). Landmark and outline methods in describing petal, sepal and labellum shapes of the flower of mokara orchid varieties. International Journal of Agriculture and Biology *13*, 652–658.

Hamilton, R.B. (1975). Comparative behavior of the American Avocet and the Black-necked Stilt (Recurvirostridae). Ornithological Monographs iii–98.

Kawabata, S., Yokoo, M., and Nii, K. (2009). Quantitative analysis of corolla shapes and petal contours in single-flower cultivars of *Lisianthus*. Scientia Horticulturae *121*, 206–212.

Kuznetsova, A., Brockhoff, P.B., and Christensen, R.H.B. (2017). lmerTest package: Tests in linear mixed effects models. Journal of Statistical Software *82*, 1–26.

Lenth, R., Singmann, H., Love, J., and others (2018). Emmeans: Estimated marginal means, aka least-squares means. R Package Version *1*.

Lindqvist, C., Motley, T.J., Jeffrey, J.J., and Albert, V.A. (2003). Cladogenesis and reticulation in the Hawaiian endemic mints (Lamiaceae). Cladistics *19*, 480–495.

Nii, K., and Kawabata, S. (2011). Assessment of the association between the three-dimensional shape of the corolla and two-dimensional shapes of petals using Fourier descriptors and principal component analysis in *Eustoma grandiflorum*. Journal of the Japanese Society for Horticultural Science *80*, 200–205.

Ortiz, P., Arista, M., and Talavera, S. (2000). Pollination and breeding system of *Putoria calabrica* (Rubiaceae), a Mediterranean dwarf shrub. Plant Biology *2*, 325–330.

Pour, A.S., Chegini, G., Zarafshan, P., and Massah, J. (2018). Curvature-based pattern recognition for cultivar classification of *Anthurium* flowers. Postharvest Biology and Technology *139*, 67–74.

Preibisch, S., Saalfeld, S., and Tomancak, P. (2009). Globally optimal stitching of tiled 3D microscopic image acquisitions. Bioinformatics *25*, 1463–1465.

Rohlf, F.J. (1990). Fitting Curves to Outlines. In Proceedings of the Michigan Morphometrics Workshop, F.J. Rolhf, and F.L. Bookstein, eds. (Ann Arbor, MI: University of Michigan Museum of Zoology), pp. 177–188.

Rohlf, F.J. (2015). The tps series of software. Hystrix *26*.

Ruan, C., Li, H., and Mopper, S. (2008). The impact of pollen tube growth on stigma lobe curvature in *Kosteletzkya virginica*: The best of both worlds. South African Journal of Botany *74*, 65–70.

Rueden, C.T., Schindelin, J., Hiner, M.C., DeZonia, B.E., Walter, A.E., Arena, E.T., and Eliceiri, K.W. (2017). ImageJ2: ImageJ for the next generation of scientific image data. BMC Bioinformatics *18*, 529–555.

Song, X., Gao, K., Fan, G., Zhao, X., Liu, Z., and Dai, S. (2018). Quantitative classification of the morphological traits of ray florets in large-flowered Chrysanthemum. HortScience *53*, 1258–1265.

Stearn, W.T. (2002). The genus *Epimedium* and other herbacious Berberidaceae including the genus *Podophyllum* (Portland, OR: Timber Press, Inc.).

Suzuki, K. (1984). Pollination system and its significance on isolation and hybridization in Japanese *Epimedium* (Berberidaceae). The Journal of Plant Research *97*, 381–396.

Webster, M., and Sheets, H.D. (2010). A practical introduction to landmark-based geometric morphometrics. The Paleontological Society Papers *16*, 163–188.

Wickham, H., Averick, M., Bryan, J., Chang, W., McGowan, L.D., François, R., Grolemund, G., Hayes, A., Henry, L., Hester, J., et al. (2019). Welcome to the tidyverse. Journal of Open Source Software *4*, 1686.
